# Comprehensive machine-learning-based analysis of microRNA-target interactions reveals variable transferability of interaction rules across species

**DOI:** 10.1101/2021.03.28.437385

**Authors:** Gilad Ben Or, Isana Veksler-Lublinsky

## Abstract

**Background:** MicroRNAs (miRNAs) are small non-coding RNAs that regulate gene expression post-transcriptionally via base-pairing with complementary sequences on messenger RNAs (mRNAs). Due to the technical challenges involved in the application of high-throughput experimental methods, datasets of direct bona-fide miRNA targets exist only for a few model organisms. Machine learning (ML) based target prediction methods were successfully trained and tested on some of these datasets. There is a need to further apply the trained models to organisms where experimental training data is unavailable. However, it is largely unknown how the features of miRNA-target interactions evolve and whether there are features that have been fixed during evolution, questioning the general applicability of these ML methods across species.

**Results:** In this paper, we examined the evolution of miRNA-target interaction rules and used data science and ML approaches to investigate whether these rules are transferable between species. We analyzed eight datasets of direct miRNA-target interactions in four organisms (human, mouse, worm, cattle). Using ML classifiers, we achieved high accuracy for intra-dataset classification and found that the most influential features of all datasets significantly overlap. To explore the relationships between datasets we measured the divergence of their miRNA seed sequences and evaluated the performance of cross-datasets classification. We showed that both measures coincide with the evolutionary distance of the compared organisms.

**Conclusions:** Our results indicate that the transferability of miRNA-targeting rules between organisms depends on several factors, the most associated factors being the composition of seed families and evolutionary distance. Furthermore, our feature importance results suggest that some miRNA-target features have been evolving while some have been fixed during evolution. Our study lays the foundation for the future developments of target prediction tools that could be applied to “non-model” organisms for which minimal experimental data is available.

Availability and implementation The code is freely available at https://github.com/gbenor/TPVOD

## Introduction

MicroRNAs (miRNAs) are small non-coding RNAs that regulate gene expression post-transcriptionally. miRNAs are encoded in the genome and are generated in a multi-stage process by endogenous protein factors [16]. The functional mature miRNAs associate with Argonaute proteins to form the core of the miRNA-induced silencing complex (miRISC). miRISC uses the sequence information in the miRNA as a guide to recognize and bind partially complementary sequences on 3’ untranslated region (UTR) of target mRNAs. miRISC binding typically leads to translational inhibition and/or degradation of targeted mRNAs [22]. miRNAs are conserved throughout evolution and are present in the genomes of animals and plants [27]. miRNAs have diverse functions in development and physiology and have been implicated in many human diseases [54].

The identification of miRNA target sites on mRNAs is a fundamental step in understanding miRNA involvement in cellular processes. Several experimental high-throughput methods for identifying miRNA targets have been developed in recent years [34,41]. The most common and straightforward approach is based on measuring changes in mRNA levels following miRNA over-expression or inhibition in tissue-cultured cells [60]. However, this approach has several major limitations [34,41]. First, such data may contain indirect signals of miRNA regulation from the downstream genes of direct miRNA targets. Second, for direct regulation, the exact sequences of binding sites are unknown and have to be predicted within the relevant mRNA sequence. Furthermore, such experimental settings may represent a non-physiological context for miRNA activity which does not reflect endogenous targeting rules. Finally, it may miss signals of translation efficiency inhibitions which consequently affect gene expression but are not reflected in changes in mRNA levels [15].

Other methods, e.g., HITS-CLIP [10,67] and PAR-CLIP [20], are based on crosslinking and immunoprecipitation (CLIP) of RNA-protein complexes that are found in direct contact. The crosslinked complexes are immunoprecipitated with a specific AGO antibody (AGO-CLIP), and the associated miRNAs and mRNA targets are collected for further sequencing analysis. Though these methods greatly decrease the target search space to precise regions on mRNAs, the identity of the specific miRNA engaged in each interaction is unknown and has to be predicted bioinformatically [61,63], by e.g., identifying which highly expressed miRNAs are associated with individual AGO-CLIP peaks [26,35,39,53].

Recently, more advanced methods, e.g., CLASH (Cross-linking, Ligation and Sequencing of Hybrids) [21], CLEAR (covalent ligation of endogenous Argonaute-bound RNAs)-CLIP [45, 55] and modified iPAR-CLIP [19], have been developed to capture miRNAs bound to their respective targets. These methods are derived from AGO-CLIP and use an extra step to covalently ligate the 3’ end of a miRNA and the 5’ end of the associated target RNA within the miRISC. Subsequent cloning and sequencing of isolated chimeric miRNA-target reads facilitate the identification of direct miRNA-target interactions. Using these methods, datasets of chimeric miRNA-target interactions were generated from cells originating from human, mouse, the nematode *Caenorhabditis elegans (C. elegans)*, and the cattle *Bos taurus*. An additional method, iCLIP [6], was applied to *C. elegans* to recover chimeric sequences without employing the ligation step. Furthermore, re-analysis of published human and mouse AGO-CLIP data discovered additional miRNA-target chimeric interactions in libraries where no ligase was added [19].

The analysis of chimeric miRNA-target interactions from the above studies revealed that many of them display non-canonical seed binding patterns and involve nucleotides outside of the seed region. Despite the great contribution that these experimental methods can bring to the miRNA field, their application is technically challenging. Thus so far, datasets were generated for only a small number of model organisms (Table 1).

**Table 1.** Datasets’ information

| Name | Cell/ Developmental stage | Experimental Method | Reference |
| --- | --- | --- | --- |
| ca1 | Madin-Darby bovine kidney (MDBK) | CLEAR-CLIP | [55] |
| ce1 | L3 staged | Modified iPAR-CLIP | [19] |
| ce2 | Mid-L4 WT (N2) | ALG-1 iCLIP endogenous ligation | [6] |
| h1 | Human embryonic kidney293 cells (HEK293) | CLASH | [21] |
| h2 | A mix of 6 datasets | AGO-CLIP endogenous ligation | [19] |
| h3 | Human hepatoma cells (Huh-7.5) | CLEAR-CLIP | [45] |
| m1 | A mix of 3 datasets | AGO-CLIP endogenous ligation | [19] |
| m2 | N2A mouse neuroblastoma (ATCC) | CLEAR-CLIP | [45] |

The limited number of experimentally identified miRNA-target interactions promoted the use of computational predictions to expand miRNA-target repertoires. Nevertheless, computational identification is very challenging, since miRNAs are short and engage only a partial sequence complementarity to their targets, and the rules that govern the miRNA targeting process are not yet fully understood. Over the past 15 years, many computational tools were developed for miRNA target prediction. The first generation of tools was based on very general rules of thumb, e.g., canonical seed pairing, miRNA-target duplex energy, conservation of the target site and accessibility [14,25,28,33]. These tools suffer from high False Positive and False Negative prediction rates [17,44,48,51], due to limitations of general rules and insufficient knowledge about seedless interactions and base pairing patterns in the non-seed region. In addition, the target prediction outputs of various tools only partially overlap, making it difficult to choose candidates for further experimental validation or more global downstream analysis.

The availability of new datasets of high-throughput, direct miRNA-target interactions (e.g., [19,21,45,55]), led to the development of new machine-learning (ML) based methods for miRNA target prediction [8,13,38,43,52,65]. These ML-based methods were designed to capture both canonical sites based on seed complementarity and non-canonical sites with pairing beyond the seed region. These methods incorporated in their models tens to hundreds of different features to represent e.g., sequence, structure, conservation, and context of the interacting molecules and were reported to achieve significant improvement in overall predictive performance than the previous tools. Differences in several aspects can be observed among ML-based methods, including the ML approach and the features used, the choice of datasets for training and testing, inclusion or exclusion of non-canonical interactions from the training/testing set, and the approach of generating negative data. We provide a summary of some of these methods, focusing on these aspects, in the supplementary material (Additional file 1, text and Table S1).

Briefly, the methods chimiRic [38] and miRTarget [36, 64] use support vector machine (SVM) for classifying miRNA-target interactions. TarPmiR [13] is a random-forest (RF) based approach that provides a probability that a candidate target site is a true target site. DeepMirTar [65], miRAW [52] and mirLSTM [49] apply deep learning approaches that are based on stacked de-noising auto-encoder (SdA), deep artificial neural networks (ANN), and long short term memory (LSTM), respectively.

In these methods, the ML models were trained and tested on a dataset of chimeric interactions from human cells generated with the CLASH method [21]. In some of the studies, the dataset was filtered based on the location of the sites, seed pairing pattern, or functional evidence. In other studies, it was complemented with additional interactions from other experiments. For example, DeepMirTar [65] and mirLSTM [49] included only canonical and non-canonical sites that are located at the 3’UTRs and added additional interactions retrieved from miRecords [66]. chimiRic [38] and miRAW [52] complemented this dataset by seed-containing sites from AGO-CLIP data, while miRTarget [64] complemented it with endogenously ligated chimeras from human AGO-CLIP experiments. miRAW [52] and miRTarget *υ4*[36] intersected the CLASH dataset with other resources to retain only interactions with functional evidence.

For additional independent testing, the above methods used few other datasets, which are not necessarily derived from ligation-based experiments. These datasets include human PAR-CLIP datasets, mouse HITS-CLIP dataset, chimeric interactions from iPAR-CLIP in *C. elegans* and CLEAR-CLIP in mouse [45], and microarray-based datasets following miRNA transfections or knockdowns (Additional file 1, Table S1).

The experimental datasets that are used to train the ML-based tools are limited to only a few model organisms. Nevertheless, there is a need to apply target prediction tools to other organisms as well, where experimental data is not available. Though some of the ML methods examined the possibility to predict interactions in organisms different from the organism they were trained on, in all cases the training was performed on human datasets and was applied on a few other organisms. The ability to do the predictions in the opposite direction, or between organisms other than human was not tested. Moreover, it is largely unknown how the patterns of miRNA-target interactions evolve across bilaterian species and whether there are features that have been fixed during evolution, thus questioning the general applicability of these ML methods across species.

In this paper, we used available datasets of high-throughput direct miRNA-target interactions to explore whether miRNA-target interaction rules are transferable from one organism to another. We evaluated eight datasets from four organisms (human, mouse, worm, and cattle), generated from various tissues and experimental protocols. We developed a processing pipeline to transform these datasets into a standard format that enables their comparison and integration. We provide a detailed overview of the datasets, focusing on their sizes, miRNA-seed families composition, and interaction patterns while highlighting their resemblance and dissimilarity. For each dataset, we trained and tested 6 commonly used machine learning classifiers for the prediction of miRNA-target interactions and evaluated the importance of the features we used. We then explored the relationships between datasets by measuring the divergence of their miRNA seed sequences, and by evaluating the performance of cross-datasets classification. Our results indicate that the transferability of miRNA-targeting rules between different organisms depends on several factors, including the composition of seed families and evolutionary distance. Our study provides important insights for future developments of target prediction tools that could be applied to organisms without sufficient experimental data.

## Results

### Dataset processing

Eight miRNA-target chimera datasets have been previously generated for human, mouse, worm (*Caenorhabditis elegans*), and cattle (*Bos taurus*). The details of each dataset are provided in Table 1, including the cell type or developmental stage that was examined and the experimental methods to obtain the data. Five of the datasets were generated by AGO-CLIP with an extra step to covalently ligate the miRNA and the target RNA (*ca1, ce1, h1, h3, m2*). An additional *C. elegans* dataset (*ce2*) contains chimeras recovered from an iCLIP experiment that did not apply an additional ligation step. Two datasets (*h2, m1*) were generated by re-analysis of published mammalian AGO-CLIP data which also recovered miRNA-chimeras in libraries where no ligase was added [19]. The *h2* and *m1* datasets contain chimeras from a mix of 6 and 3 independent experiments, respectively.

We applied a multi-step processing pipeline (step 1) to retrieve all the required information about the interactions (e.g., miRNA name and sequence, target site sequence, and location). Since miRNA target sites that are located at the 3’UTRs of mRNA sequences are considered to be most functional [3,43], (step 2) we filtered out non 3’UTR interactions. Then (step 3) we generated miRNA-target duplexes using RNAduplex [37]. Finally (step 4) we classified the interactions based on their seed-pairing patterns and kept only interactions with canonical or non-canonical pairing patterns (see definitions in methods).

The numbers of interactions that passed the pipeline stages are shown in Table 2. The 3’UTR interactions represent 10%-47% of all interactions in the datasets and interactions with canonical or non-canonical seed-pairing constitute 53%-82% of them. The pipeline produced final datasets with variable sizes: four small datasets (500-1200), two large datasets (2000-5000), and two massive datasets (~18,000 each). These final datasets were later used as input for machine learning tasks. Therefore, we complemented the datasets with synthetically-generated negative interactions as described in methods. We extracted 490 features from each interaction, representing properties of the interaction duplex and interaction site and its flanking region within the 3’UTR (see methods: Features).

**Table 2.** Summary of the data processing pipeline

| Dataset | ca1 | ce1 | ce2 | h1 | h2 | h3 | m1 | m2 |
| --- | --- | --- | --- | --- | --- | --- | --- | --- |
| No. of interactions <sup>a</sup> | 296,297 | 3,627 | 4,920 | 18,514 | 10,567 | 32,712 | 1,986 | 130,094 |
| No. of interactions in 3'UTRs | 30,534 | 1,704 | 1,206 | 8,507 | 2,039 | 4,634 | 902 | 33,100 |
| **Final dataset (canonical & non-canonical interactions)** | **18,204** | **1,176** | **992** | **5,137** | **1,150** | **2,846** | **537** | **17,574** |

<sup>a</sup> As provided by the original publications

### Datasets’ characteristics

In the following subsections we characterized the interactions of each dataset based on their miRNA distribution and base-pairing patterns. Since the negative interactions are synthetically-generated, we focused on positive interactions only.

#### miRNA distribution

We counted the appearance of miRNA sequences and miRNA seed families (nt2-7) and generated a distribution function for each dataset (Table 3). Our analysis indicates that the datasets are not uniformly distributed in terms of miRNA appearances (Figure 1). Furthermore, 90% of the interactions are dominated by a small subset of miRNA sequences (25-50%) or miRNA seed families (18-37%).

**Figure 1.**
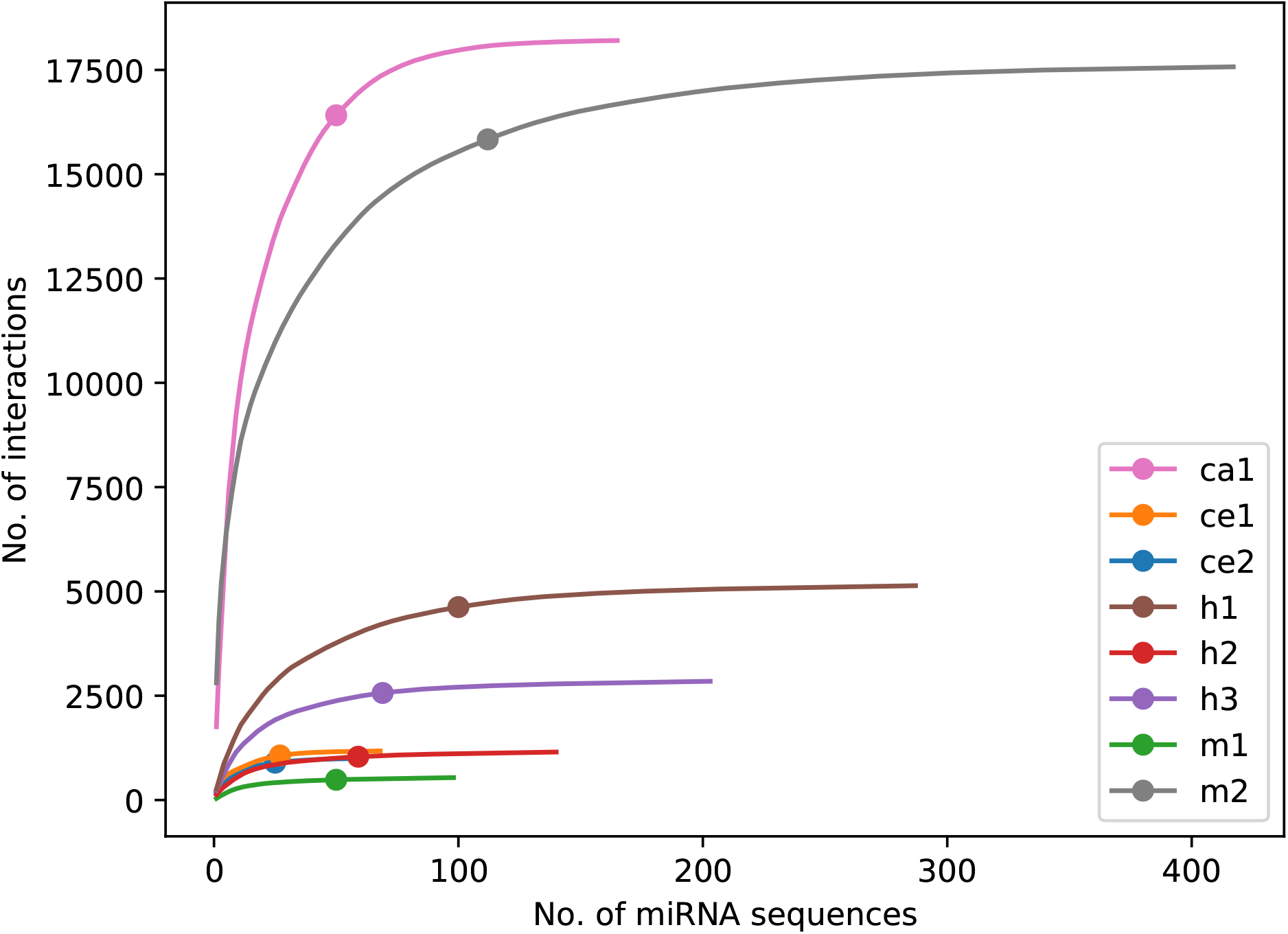
Cumulative sum of miRNA sequence appearances in the examined datasets. Each curve corresponds to the cumulative sum of one of the datasets, where the bold points indicate the minimum number of unique miRNA sequences needed to represent 90% of the interactions within a dataset. The height of a curve represents the size of the dataset, and the width of a curve represents the number of unique miRNA sequences that comprise the dataset.

**Table 3.**
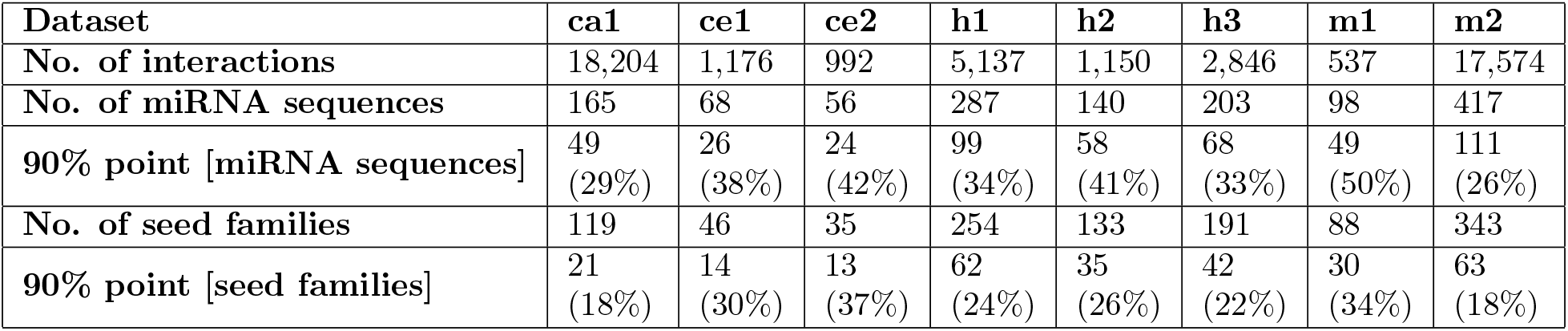
Composition of miRNA sequences and miRNA seed families within datasets

#### Seed types and base-pairing density

We classified the interactions (i.e., the corresponding duplexes formed by the miRNA and the target site) based on two parameters: seed type (canonical or non-canonical, see methods) and base-pairing density (number of base-pairs (bp) within the duplex: low with less than 11bp, medium with 11-16bp and high with more than 16bp). We defined 6 classes based on combinations of seed type and base-pairing density and assigned each interaction to the appropriate class (Figure 2). As can be seen in the figure, the datasets are rich and diverse and include all the combinations of seed type and base-pairing density. Nevertheless, two observations stand out: (1) In terms of seed type, the majority of the interactions are non-canonical (48-70%); and (2) for both classes of seed types, the majority of the interactions have medium and high base-pairing density, while the low base-pairing density interactions comprise only a small portion of the datasets. Similar analysis for the negative interactions is shown in Additional file 1, Figure S1.

**Figure 2.**
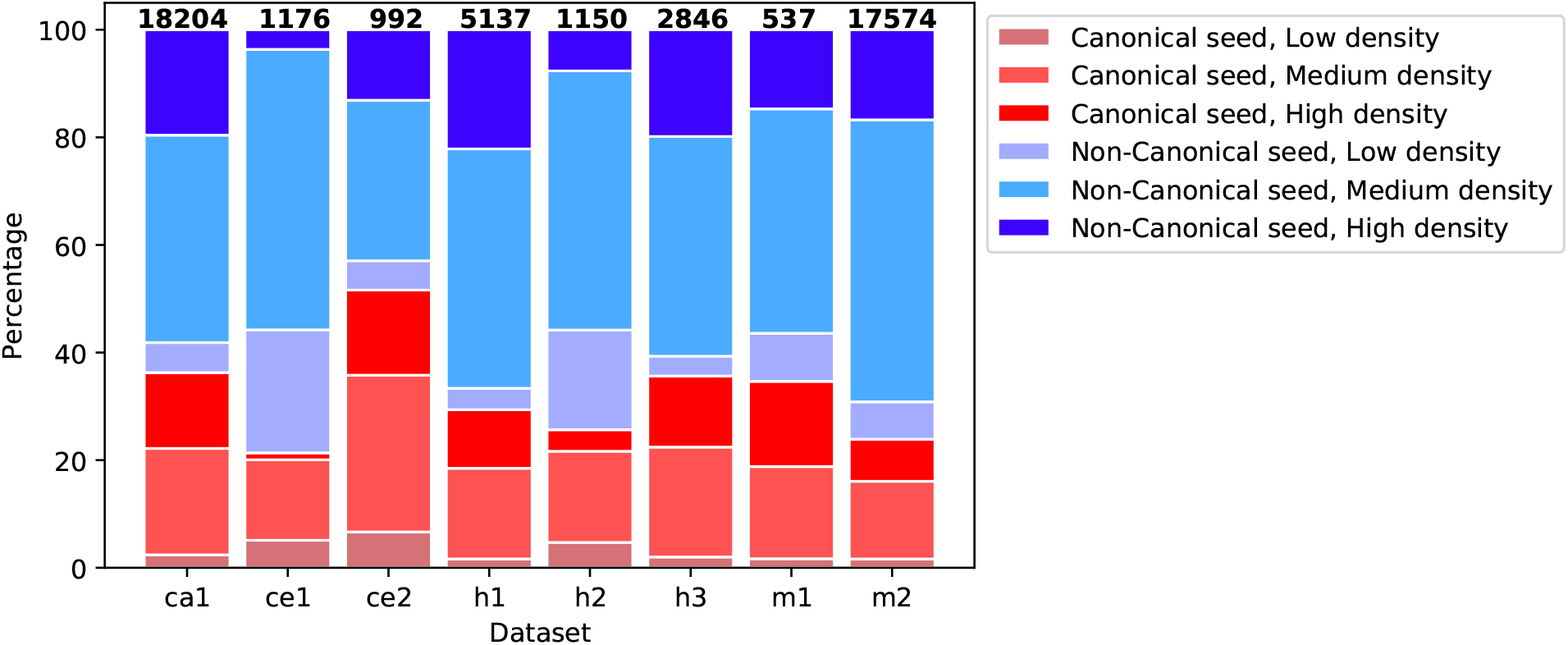
Classification of the miRNA-target duplexes based on their base-pairing patterns. Distribution of miRNA-target duplexes across 6 classes according to the seed type (canonical and non-canonical) and to the base-pairing density (low: less than 11bp, medium: 11-16bp and high: more than 16bp).

### Intra-dataset analysis

In this section, we evaluated the performance of machine-learning-based binary classifiers to correctly classify positive and negative miRNA-target interactions within the same dataset. We first conducted a set of experiments with different types of commonly used machine learning classifiers. Then, we performed an in-depth analysis of the best performing classifier, by measuring different performance metrics and by estimating feature importance.

#### Evaluation of different machine learning methods

For each dataset, we generated 20 training-testing splits of the data using a stratified random split algorithm. This split algorithm ensures that each miRNA appears in the training and the testing sets at the same proportion as in the original dataset. We then trained 6 widely used classifiers on the 20 training sets of each dataset and measured their performance in the classification of their respective testing sets. We calculated the mean and the standard deviation values of the classification accuracy as shown in Table 4. Notably, XGBoost classifier achieved the best results across all datasets, with accuracy scores ranging from 0.82 to 0.94, with the following order of performance from low to high: *h1, h3, m1, ce1, ce2, m2, h2, ca1*. We did not observe any bias in the ordering of the organisms in the list. We compared our results to previous machine learning-based approaches that were trained and tested on the human CLASH dataset designated as *h1* in Table 1. The accuracy achieved by our classifiers on this dataset is comparable to the ones reported by previous studies (Additional file 1, Table S2).

**Table 4.** Intra-dataset classification accuracy of different machine learning methods

| Dataset | XGBoost | RF | KNN | SGD | SVM | LR |
| --- | --- | --- | --- | --- | --- | --- |
| ca1 | 0.937<br>(0.002) | 0.885<br>(0.004) | 0.828<br>(0.003) | 0.797<br>(0.033) | 0.895<br>(0.003) | 0.836<br>(0.004) |
| ce1 | 0.889<br>(0.014) | 0.833<br>(0.019) | 0.768<br>(0.019) | 0.798<br>(0.045) | 0.841<br>(0.015) | 0.843<br>(0.014) |
| ce2 | 0.891<br>(0.016) | 0.858<br>(0.018) | 0.768<br>(0.019) | 0.819<br>(0.034) | 0.862<br>(0.012) | 0.847<br>(0.016) |
| h1 | 0.824<br>(0.007) | 0.769<br>(0.008) | 0.731<br>(0.007) | 0.746<br>(0.011) | 0.795<br>(0.007) | 0.770<br>(0.007) |
| h2 | 0.904<br>(0.007) | 0.869<br>(0.011) | 0.857<br>(0.009) | 0.860<br>(0.03) | 0.879<br>(0.009) | 0.892<br>(0.009) |
| h3 | 0.835<br>(0.007) | 0.769<br>(0.009) | 0.744<br>(0.009) | 0.752<br>(0.034) | 0.805<br>(0.007) | 0.795<br>(0.010) |
| m1 | 0.847<br>(0.015) | 0.795<br>(0.016) | 0.758<br>(0.022) | 0.760<br>(0.038) | 0.819<br>(0.019) | 0.800<br>(0.019) |
| m2 | 0.900<br>(0.004) | 0.826<br>(0.004) | 0.797<br>(0.004) | 0.798<br>(0.017) | 0.873<br>(0.004) | 0.833<br>(0.004) |

The cells contain the mean and the standard deviation (in brackets) values of the accuracy results acquired from 20 models that were trained and evaluated on different training-testing dataset splits

#### In-depth analysis of the XGBoost performance

We next sought to perform an in-depth performance analysis for the XGBoost classifier since it achieved the highest accuracy across all datasets. Therefore, we calculated 5 additional commonly used performance metrics. The area under the curve (AUC) is a performance measurement for classification problems at different threshold settings. It provides information on the capability of a model to differentiate between classes. AUC values range from 0 to 1, where a model with perfect predictions achieves AUC of 1. True positive rate (TPR) and true negative rate (TNR) are the percentages of actual positive or negative results that are correctly identified. For ideal classifiers, these metrics are close to 1. The Matthews correlation coefficient (MCC) is used as a measure of the quality of classifications. A coefficient of +1 represents a perfect prediction, 0 an average random prediction, and −1 an inverse prediction. The F1 score is an average of two metrics: precision (proportion of positive classification which was correct) and recall (proportion of positives that were correctly classified). The F1 score reaches its best value at 1 and its worst score at 0.

We observed that the values for all datasets are similar and relatively high. Table 5 summarises the performance metrics of the XGBoost classifiers for each dataset. As before, we calculated the mean scores and the standard deviation values of the metrics across 20 training-testing data splits. The average metrics scores ranged as follows: AUC score ranged in 0.91-0.98, TPR and TNR ranged in 0.82-0.91, MCC coefficient ranged in 0.65-0.87, and F1 score ranged in 0.82-0.94. In accordance with the accuracy metric previously calculated, these metrics indicate that all 8 XGBoost classifiers (corresponding to each dataset) are accurate, balanced, and precise.

**Table 5.** XGBoost performance measurements

| Dataset | AUC <sup>a</sup> | ACC <sup>b</sup> | TPR <sup>c</sup> | TNR <sup>d</sup> | MCC <sup>e</sup> | F1 score |
| --- | --- | --- | --- | --- | --- | --- |
| ca1 | 0.983<br>(0.001) | 0.937<br>(0.002) | 0.932<br>(0.004) | 0.943<br>(0.004) | 0.874<br>(0.004) | 0.937<br>(0.002) |
| ce1 | 0.955<br>(0.009) | 0.889<br>(0.014) | 0.89<br>(0.018) | 0.889<br>(0.014) | 0.779<br>(0.028) | 0.89<br>(0.014) |
| ce2 | 0.958<br>(0.012) | 0.891<br>(0.016) | 0.884<br>(0.02) | 0.899<br>(0.019) | 0.783<br>(0.032) | 0.89<br>(0.017) |
| h1 | 0.908<br>(0.006) | 0.824<br>(0.007) | 0.816<br>(0.008) | 0.833<br>(0.008) | 0.649<br>(0.014) | 0.822<br>(0.007) |
| h2 | 0.972<br>(0.003) | 0.904<br>(0.007) | 0.886<br>(0.012) | 0.924<br>(0.011) | 0.809<br>(0.014) | 0.902<br>(0.007) |
| h3 | 0.914<br>(0.004) | 0.835<br>(0.007) | 0.823<br>(0.011) | 0.849<br>(0.009) | 0.671<br>(0.014) | 0.832<br>(0.008) |
| m1 | 0.914<br>(0.007) | 0.847<br>(0.015) | 0.834<br>(0.014) | 0.862<br>(0.024) | 0.695<br>(0.031) | 0.844<br>(0.014) |
| m2 | 0.963<br>(0.002) | 0.9<br>(0.004) | 0.891<br>(0.003) | 0.909<br>(0.005) | 0.8<br>(0.008) | 0.899<br>(0.004) |

<sup>a</sup> Area Under the Receiver Operating Characteristic Curve
<sup>b</sup> Overall accuracy
<sup>c</sup> True Positive Rate (Sensitivity)
<sup>d</sup> True Negative Rate (Specificity)
<sup>e</sup> Matthews correlation coefficient
The cells contain the mean scores and the standard deviation (in brackets) values acquired from 20 models that were trained and evaluated on different training-testing dataset splits.

#### Random split control

In the above analysis, the splitting of the dataset into the training and testing sets was done in a way that preserves the miRNA distribution in both sets (stratified split). To assess how this type of split affects the classifier performance, we repeated the analysis with the XGBoost classifier, but this time used a random split strategy (control split) to generate the training and the testing sets. Our results showed that there is almost no difference between the results achieved with the stratified and the control split methods. The accuracy results for the control splits are found in Additional file 1, Table S3.

#### Top important features of each dataset

We used 490 features to describe the interactions. We next sought to identify the top important features of each dataset, their relative scores, and the degree of overlap of the top features between different datasets. XGBoost classifier reports a list of five feature importance metrics: weight, gain, cover, total gain, and total cover. We extracted the 5 metrics for all the 20 training-testing splits of each dataset and calculated their mean and standard deviation (Additional file 2, Table S8). For further analysis, we chose the gain metric, which reflects the contribution of each feature to the model. For each dataset, we sorted the features in descending order, based on their mean gain score. The plots of the feature importance curves for all datasets are shown in Figure 3. Figure 3a reveals that the gain score is decaying very fast. The 6 top features are significantly stronger relative to the rest (Figure 3b). We thus extracted the top 6 features from each dataset, along with their scaled gain score (see methods), into a unified list. The unified list consisted of a total of 16 features (out of the maximum length of 48), indicating that there are many shared features among the datasets. Table 6 shows the features ordered by their mean gain across all datasets and the top six features of each dataset are marked with a star. As can be seen in the table, there are at least 3 features common to each dataset pair. In addition, only a small number of features belong to a single dataset, indicating that the features in the unified list may well represent all 8 datasets. Notably, features related to the seed region (marked as bold in the table), comprise half of the features in the list. This finding emphasizes the role of the seed region in the formation of miRNA-mRNA interactions.

**Figure 3.**
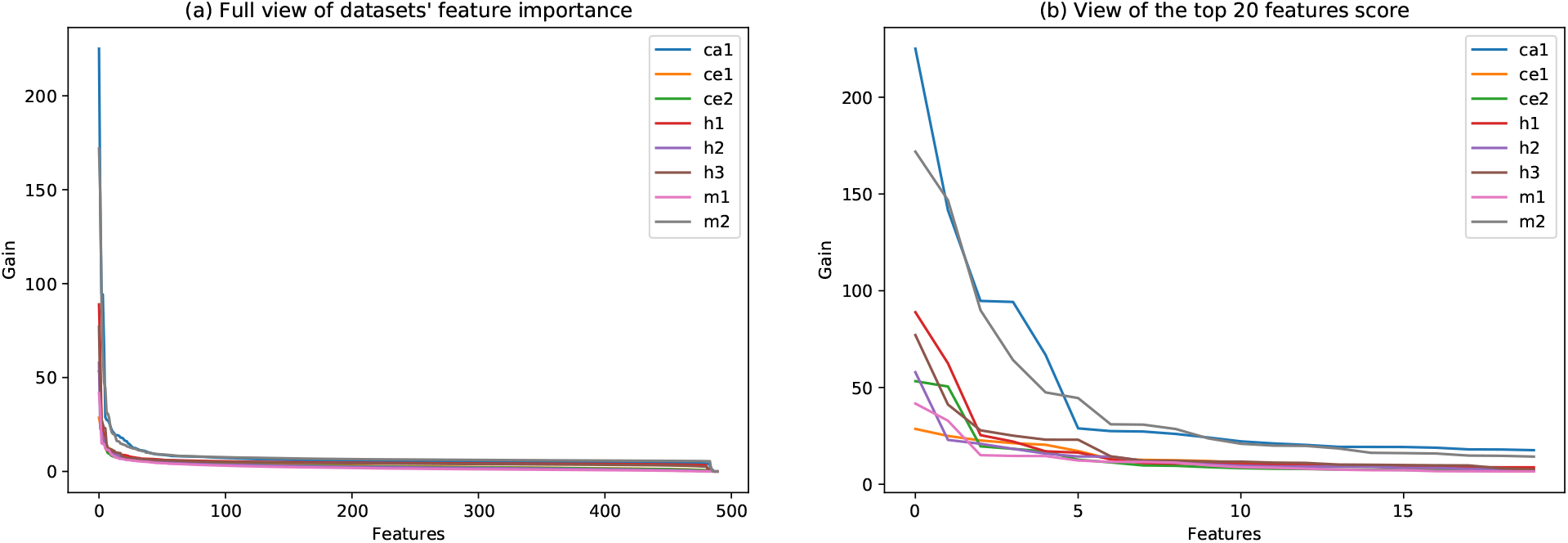
Datasets’ feature importance plot based on gain score. The features are sorted in descending order from the top feature (highest gain) to lowest. (a) A full view of the gain plot emphasizes the gain decay. (b) A zoomed view, focused on the top 20 features.

**Table 6.** Feature importance

| Feature/Dataset | ca1 | ce1 | ce2 | h1 | h2 | h3 | m1 | m2 | mean |
| --- | --- | --- | --- | --- | --- | --- | --- | --- | --- |
| Number of GU bp within the seed<sup>a</sup> | 100* | 87* | 95* | 29* | 40* | 100* | 28 | 100* | 72 |
| bp in the 1st nt of the seed<sup>b</sup> | 63* | 79* | 34* | 70* | 25* | 30* | 27 | 85* | 52 |
| Number of GU bp within the site<sup>a</sup> | 42* | 71* | 32* | 100* | 19 | 53* | 35* | 28* | 48 |
| Proportion of G in mRNA at the site region<sup>a</sup> | 12 | 74* | 12 | 12 | 36* | 33* | 100* | 37* | 39 |
| Duplex minimum free energy<sup>a</sup> | 13* | 45 | 11 | 10 | 100* | 19 | 35* | 52* | 36 |
| Number of bp at location 2-7<sup>a</sup> | 42* | 33 | 100* | 12 | 18 | 36* | 13 | 18 | 34 |
| Proportion of GG in mRNA at the site region<sup>a</sup> | 30* | 21 | 10 | 12 | 7 | 30* | 79* | 26* | 27 |
| bp in the 4th nt of the seed<sup>b</sup> | 8 | 100* | 21 | 10 | 11 | 16 | 2 | 12 | 22 |
| Number of bulges outside the seed<sup>a</sup> | 3 | 60* | 6 | 25* | 32* | 9 | 9 | 8 | 19 |
| bp in the 2nd nt of the seed<sup>b</sup> | 8 | 42 | 37* | 7 | 11 | 13 | 15 | 6 | 17 |
| bp in the 5th nt of the seed<sup>b</sup> | 12 | 27 | 14 | 14 | 6 | 15 | 29* | 12 | 16 |
| Number of GC bp within the seed<sup>a</sup> | 7 | 22 | 24* | 18* | 12 | 13 | 11 | 12 | 15 |
| Number of GC bp outside the seed<sup>a</sup> | 4 | 27 | 11 | 10 | 27* | 8 | 6 | 5 | 12 |
| Accessibility (nt=21, len=10)<sup>a</sup> | 9 | 19 | 7 | 6 | 25 | 7 | 12 | 7 | 11 |
| minimum free energy of the target site + 50nt flanking regions<sup>a</sup> | 8 | 11 | 6 | 7 | 8 | 11 | 36* | 6 | 11 |
| Number of mismatches inside the seed<sup>a</sup> | 4 | 3 | 15 | 19* | 0 | 13 | 2 | 9 | 8 |

\* Belongs to the top 6 features of the dataset
<sup>b</sup> Boolean feature
<sup>a</sup> Numeric feature

### Cross-dataset analysis

In the previous section, we trained, optimized, and evaluated the performance of a dedicated classifier for each dataset. Next, we examined the relationships between datasets. To that end, we first used a statistical measure to calculate the distance between the datasets. Second, we visualized the datasets based on the unified list of the 16 most important features that were found above. Finally, we evaluated the performance of each dataset specific classifier to properly classify interactions in the other datasets.

#### Kullback–Leibler divergence

We hypothesized that a pair of datasets, with similar characteristics, might achieve better results in classifying the interactions of each other. We thus looked for a measure to assess the level of similarity of a pair of datasets, that will take into account the directionality of the classification task: classifier is trained on one dataset (source) and is applied to classify a second dataset (target). We chose the asymmetric measure Kullback–Leibler (KL) divergence which measures how the target’s probability distribution is different from the probability distribution of the source. KL divergence has its origins in information theory which deals with the quantification of the amount of information in a given data. KL divergence is widely used to assess the approximation of samples that come from one distribution by samples that come from another distribution. The KL divergence measures the information loss in the approximation. It is typically used to measure the information loss in a case when a simpler distribution (such as a Uniform or Gaussian distribution) approximates experimental data.

Similarly, we used the KL divergence to measure the pairwise information loss between every two datasets which will later be used as training and testing sets in the analysis described below. For each pair, we calculated the divergence that can be interpreted as the amount of information lost when the training set represents the testing set. The calculation is done based on the datasets’ miRNA seed family distribution (see methods).

Figure 4 shows the divergence between all dataset pairs. The divergence of a dataset with itself is zero, and the divergence scores between datasets within the same organisms are usually lower than the divergence scores between different organisms. Notably, the divergence scores between *C. elegans* datasets, both as targets and as sources, and the other datasets are significantly higher (range in 5-8.1) than the divergence scores between other pairs (range in 1.2-3.8), indicating that seed distributions of other organisms poorly represent the *C. elegans* datasets and vice versa. The asymmetry of the KL measure can be observed, for example, for the pair *(h1,h3)*, for which *KL(target=h3 || source=h1)=1.6* and the *KL(h1 || h3)=2.1*.

**Figure 4.**
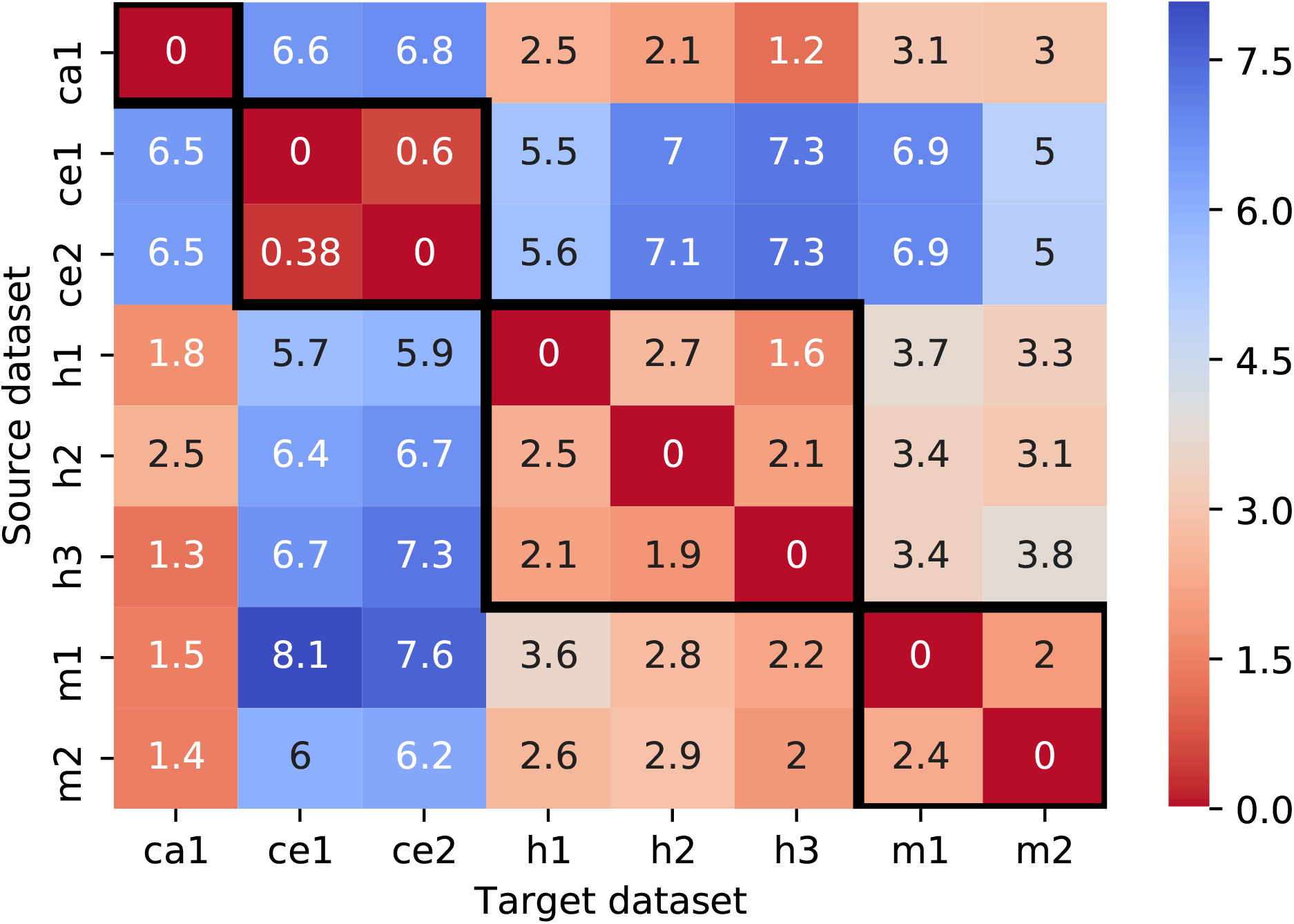
Kullback–Leibler (KL) divergence of all dataset pairs. Each cell (*i,j*) represents the divergence from a source datasets *i* to a target dataset *j* (KL(j || i)), based on their miRNA seed family distributions. The black frames surround the results of dataset pairs originating from the same organism.

Intuitively, this means that dataset *h1* better approximates dataset *h3* and there is less information loss than in the vice-versa case.

#### Dataset visualization

Visualization is an important step in the analysis of high-throughput biological data and can assist in revealing hidden phenomena. However, visualization is challenging when the data is represented by a large number of features. The dimensionality reduction algorithm enables the representation of the data in a 2-dimension scatter plot and facilitates the inspection of the data visually. To visualize the datasets in two dimensions, we first extracted for each dataset the unified list of 16 top features found above (see Table 6) and then performed a dimensional reduction using the PCA technique. The results are shown in Figure 5.

**Figure 5.**
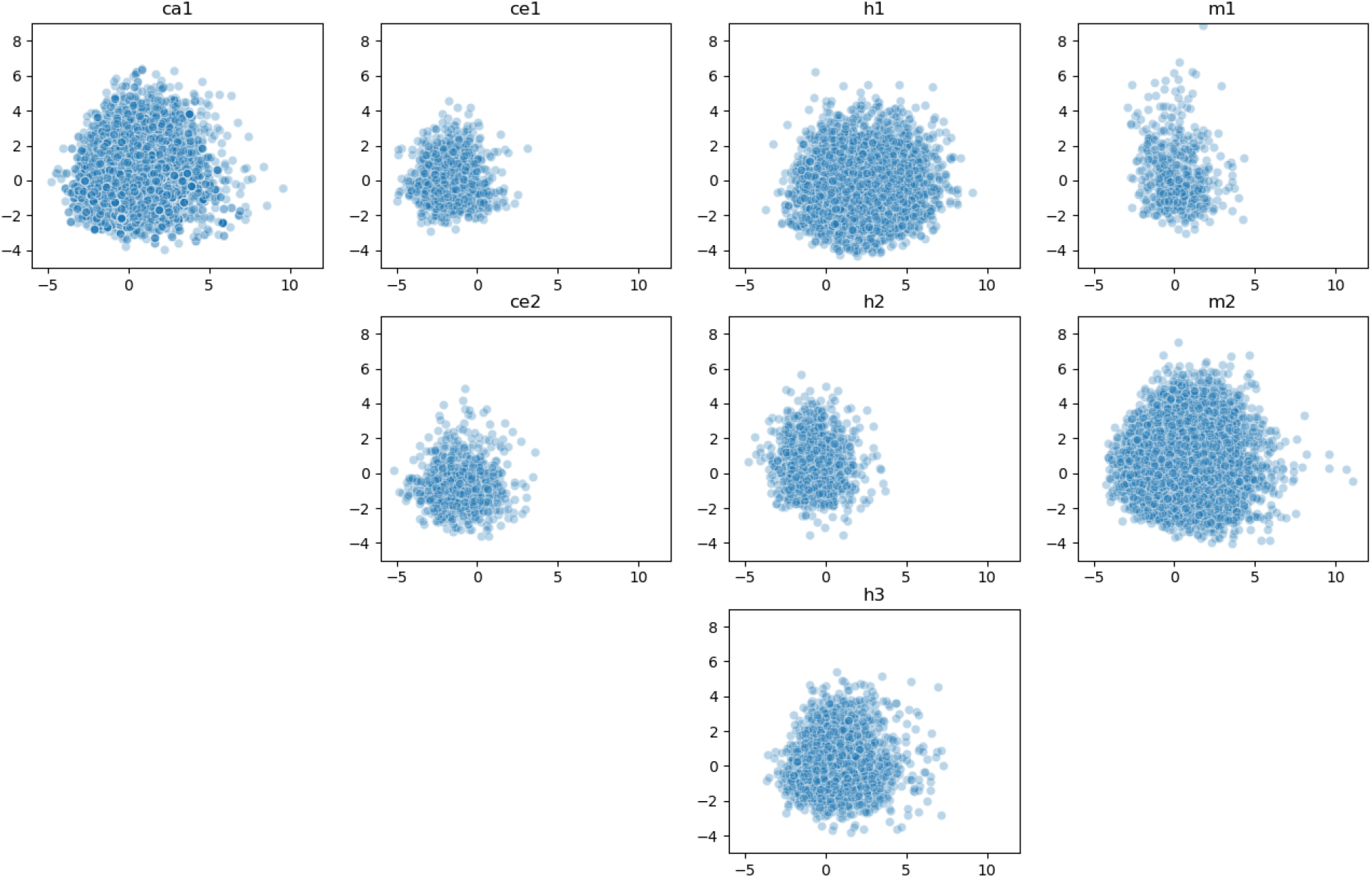
Visualization of the datasets in 2D. Each point represents a single interaction after a dimensional reduction of its features’ space using PCA. X and Y axes are the first and the second components of the PCA, respectively.

Figure 5 reiterates the fact that there are big differences in the sizes of the datasets, reflected in the density of the graphs. For example, the size of the human dataset *h1* is more than twice the size of the datasets *h2* and *h3*, and indeed, its graph is denser. In addition, there are notable differences in the 2-dimensional space spanned by each dataset: while the datasets *ca1, h1, h3, m2* are spread in the whole area, *C. elegans* datasets (*ce1, ce2*) and datasets composed of endogenously ligated chimeras from a mixture of experiments (*h2, m1*) are concentrated in a narrower part of the area.

#### Classification performance between datasets

We evaluated the performance of cross dataset miRNA-target predictions, i.e., the performance of a classifier when applied to interactions from datasets different from the one it was trained on. We examined all possible 56 combinations, considering each dataset both as a training and as a testing set. For each dataset, we loaded the 20 XGBoost classifiers that we trained in section Intra-dataset analysis and used them to classify the rest 7 datasets. Figure 6 shows for every pair of datasets the mean classification accuracy over the 20 tests (for standard deviation values, see Additional file 1, Table S6.)

**Figure 6.**
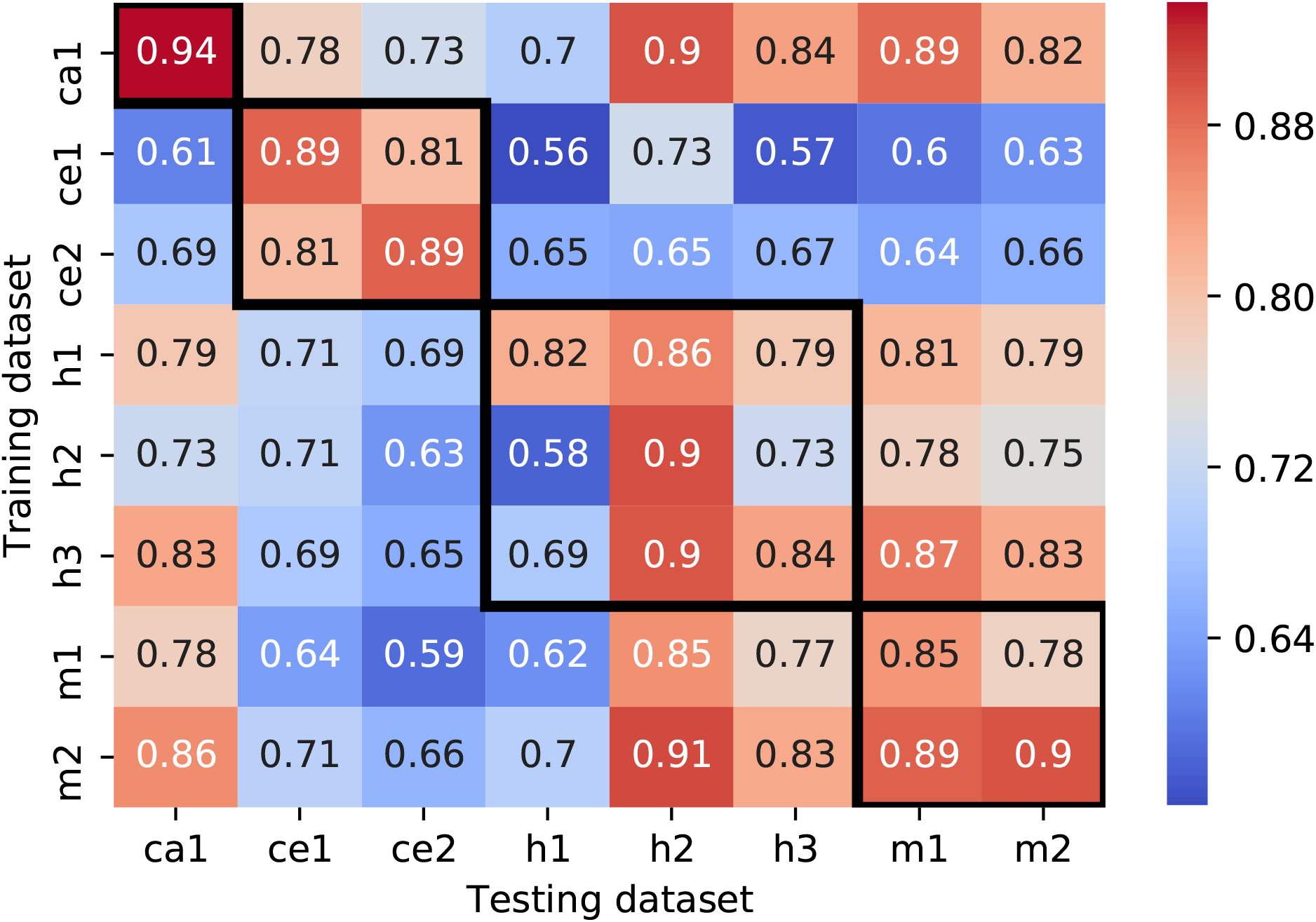
Cross-dataset classification results. Each cell *(i,j)* represents the mean accuracy of the 20 classifiers that were trained on dataset *i* (in section Evaluation of different machine learning methods) and tested on dataset *j* (ACC(i, j)). The black frames surround the results of dataset pairs originating from the same organism. The accuracy results for pairs *(i,i)*, were taken from section Intra-dataset analysis. Note: The color scale of this figure is inverse to the scale used for KL-divergence plot in Figure 4

Inspection of the results (excluding the diagonal) reveals that there is a variability in the classification performance among the pairs, ranging from random, slightly above 0.5, to 0.91. The accuracy matrix is not symmetric, i.e., a pair where a dataset *i* serves as a training set and a dataset *j* serves as a testing set, achieves a different performance than a swapped pair. Pairs of datasets originating from the same organism (surrounded by black boxes in Figure 6) generally achieved higher accuracy than pairs from mixed organisms. Intriguingly, the human pairs (*h2,h1*), (*h2,h3*), (*h3,h1*) achieved a relatively low accuracy score. The low performance of these pairs could be potentially explained by the differences in the diversity of the datasets. In particular, the dataset *h2* is smaller and less diverse then the datasets *h1,h3* (Figure 5), and thus a model that uses it as a training set achieves lower performance. In most of the cases, the KL divergence results coincide with the accuracy results. For example, for the pair (*h1,h3*), the *KL(h3* || *h1)*=1.6 <*KL(h3* || *h1)*=2.1 while *ACC(train=h1, test=h3)*=0.79 > *ACC(h3,h1)*=0.69, demonstrating that the dataset *h1* better represents the dataset *h3* and as such achieved better accuracy results than vice versa. Interestingly, the *KL(h2* || *h1)*=2.5≈*KL(h1* || *h2)*=2.7 however the *ACC(h1,h2)*=0.86 > *ACC(h2,h1)*=0.58. This indicates that there are additional factors that affect the ability to accurately classify miRNA-target interactions, such as the patterns of interactions that appear in them.

Pairs of datasets originating from different organisms, that included *C. elegans* as either a training or a testing set achieved poor performance, which ranged from 0.56 to 0.78. As we previously saw, the divergence scores of these pairs are 2x-4x larger (range from 5 to 8) than the scores of the other pairs. This may indicate that the seed distributions of human, mouse and cattle datasets are not well represented by the seed distributions of *C. elegans* datasets and vice versa. Other pairs of two organisms achieve much higher accuracy, reaching up to 0.91. The lowest accuracy in these mixed pairs was observed for pairs that contained *h1* as a testing set. Notably, this dataset was used by previous methods (Additional file 1, Table S1) for training/testing purposes only, and has never been evaluated as an independent testing set. Additional factors that could influence the classification accuracy will be further discussed in the discussion section.

## Discussion

Identification of bona-fide miRNA targets is crucial for elucidating the functional roles of miRNAs and remains a major challenge in the field. Notable progress in this task has been achieved due to novel experimental protocols that can produce high-throughput unambiguous interacting miRNA-target datasets. However, due to technical challenges involved in the application of these methods, there is a constantly increasing interest in using computational approaches for miRNA target prediction, especially those that are based on advanced machine learning models. Several studies successfully trained and applied classic machine learning [13, 36, 38, 64] and deep learning [49, 52, 65] methods on some of the experimental miRNA-target datasets from a few model organisms. However, our limited understating of the evolution of miRNA-target interactions, puts in question the applicability of these tools to organisms with no available experimental training data.

The ultimate goal of this study was to evaluate the transferability of miRNA-target rules between the examined organisms, as well as to identify and compare their most influential interaction features. To this end, we systematically characterized the available miRNA-target chimeric datasets and conducted intra- and cross-datasets classification analyses using machine learning approaches.

### Available data

The availability of large and high-quality data is crucial for machine learning-based research. In the field of experimental miRNA-target identification, several approaches to generate high-throughput datasets are available, each one has its benefits and limitations [34, 41].

In our analysis, we focused on chimeric miRNA-target datasets, generated by experimental or endogenous ligation (by e.g., CLASH [21] or PAR-CLIP [19]), as these datasets provide direct evidence for interactions between a miRNA and a specific target site. Furthermore, these datasets contain many non-canonical interactions that enrich the repertoire of miRNA-target interactions. On the other hand, the main limitation of ligation-based methods is the low yield of chimeric reads that are being recovered (around ~ 2%), suggesting that a large number of miRNA-target interactions remains uncaptured. In this work, we assume that the captured interactions represent an unbiased sampling of all the interactions in the examined cells. Additional advances in the efficiency of ligation-based methods and deeper sequencing will provide richer datasets that would be easily incorporated into our analysis for further research.

We utilized eight available chimeric datasets from four organisms (worm to human), that were generated by different experimental protocols (Table 1). We developed a processing pipeline to transform and unify the different data formats that we encountered during the collection of the datasets. This pipeline is a powerful infrastructure that will enable us, with a relatively low effort, to add more data sources to the analysis in the future, when these become available.

### Thorough analysis of the datasets

We characterized the datasets based on their miRNA content and base-pairing patterns. Our analysis of the frequencies of miRNA sequences revealed that there are differences in miRNA sequence distributions among datasets, even if they originated from the same organism. In addition, each dataset is dominated by a small set of miRNAs (30-50% of most frequent miRNAs comprise 90% of the interactions). These distributions mirror the in vivo distributions, as it was reported that miRNA frequency in miRNA-target chimeras is correlated with total miRNA abundance [45].

We continued categorizing the interactions based on their seed-pairing type (canonical and non-canonical) and base-pairing density. Perfect seed complementarity (referred to as canonical seed pairing) between target sites and miRNA seed sequences (nts 2-7 or 2-8), has long been recognized as a critical dominant feature which determines miRNA targeting efficiency [4, 33, 56]. Nevertheless, in recent years several examples of functional miRNA-target interactions without perfect seed pairing have been reported, featuring *GU* pairs, mismatches, and bulges in the seed region (referred to as non-canonical seed pairing). Examples include the well-established *let-7* targeting of *lin-41* in *C. elegans* [57,62], with one site containing one-nucleotide bulge in the target, and the other site containing a *GU* pair. Moreover, non-canonical miRNA-target sites known as “nucleation bulges”, in which the target sites contain a bulged-out *G* in the seed, were identified for *miR-124* when analyzing AGO HITS-CLIP data from mouse brain [9]. The functionality of non-canonical sites is still a matter of debate. While studies that generated miRNA-target chimeras provided evidence for the functionality of the recovered non-canonical interactions [19,21], a recent analysis of non-canonical target sites revealed that even though these sites are bound by the miRNA complex, they do not appear to be broadly involved in the regulation of gene expression [1]. Future work will need to focus on generating miRNA functional high-throughput datasets [59] across organisms, that would be combined with datasets of chimeric interactions, to provide a more robust starting point for similar types of studies.

We showed that the majority of the interactions in most datasets are non-canonical (48-70%). Furthermore, in both canonical and non-canonical groups, a large fraction of the interactions is characterized by a medium and a high density of base-pairing (11-16 and more than 16 base-pairs, respectively), predicting the existence of additional pairing beyond the seed region. These auxiliary non-seed interactions were suggested to compensate for imperfect seed matches [5, 18]. Moreover, non-seed interactions were also shown to contribute to target specificity among miRNA seed family members (same seed, divergent non-seed sequence), both in the case of canonical and non-canonical seed pairings [6, 45].

### Features and their significance

In this work, we partially adapted the pipeline from DeepMirTar [65]. In DeepMirTar the interactions are represented by 750 features. These features include high-level and low-level expert-designed features that represent the interacting duplex, sequence composition, free energy, site accessibility and conservation. Additional raw-data-level features encode the sequences of the miRNA and the target site. We have adopted some of the expert-designed features in our study and used a total of 490 various features to describe the interactions, enabling the model to identify and learn different interaction patterns.

We did not include, however, the previously suggested raw-data-level features, to avoid potential information-leakage from the training set to the testing set. First, we saw that the miRNA seed families are not uniformly distributed. Second, in our study, the negative sequences are synthetically generated such that there is no match in the seed region to any annotated miRNA. Thus, including these features could lead the machine learning model to learn to distinguish between real and mock miRNA seeds. Moreover, in such a case, the model may be over-fitted and fail to generalize the rules of interactions. Indeed, and perhaps not surprisingly, we achieved higher classification performance when we included the raw-data-level features in our models (Additional file 1, Table S4). Another study [52] that used raw sequence features addressed this issue by generating a negative dataset based on experimentally verified data instead of using mock miRNAs. A comparison between different methods for the generation of negative datasets is an interesting direction for future research. In particular, the evaluation of how the combination of these methods and different feature sets affects the performance of miRNA-target prediction classifiers would help to generate standard approaches for future studies.

The feature importance analysis revealed that there is a small group of significantly dominant features in all datasets. Even though the analysis identified the features for each dataset independently, we showed that there is a significant overlap between the groups, and the unified group contains only 16 features (Table 6). Importantly, half of these features are seed-related, reiterating the significance of this region in miRNA-target interactions [1].

Ideally, in machine learning, we want the ratio between samples and features to be high enough to have a robust model and avoid over-fitting. Some of the datasets in our collection are relatively small, with a low ratio of interactions to features. For *ce1, ce2, h2* the ratio is ~4, and for *m1* it is ~2. Low ratio can produce models with high bias and high variance. In general, a reduction in the number of features, when possible, was shown to be a successful practice. In this work, some of the features are highly correlated, thus can be combined. There are several methods for feature selection and dimensionality reduction that may be evaluated in the future. As a preview, we used a basic method for feature selection, based on the XGBoost feature importance data. We used the 16 features taken from Table 6 and repeated the classification analysis (Additional file 1, Table S7, Figure S2). The results were similar to the results obtained when all features were included, indicating that future research that will evaluate different dimensionality reduction methods should be considered to optimize the classification models.

### Training and testing dataset split

The splitting procedure of the data into training and testing sets has a crucial role in the evaluation of machine learning models. In the miRNA-target prediction task, there is no pre-defined split to training and testing sets as is usually common in other fields, for example, in computer vision (e.g., MNIST [30]). Therefore, we used three strategies to reduce the effect of the split on our results: (1) using stratified training-testing split which ensures the same distribution of miRNA sequences in both training and testing sets; (2) generating control sets by a pure-random split algorithm and (3) generating for the above split approaches several training-testing sets using different random states and reporting the mean and the standard deviation values of the results. Indeed, we got similar results with both splitting methods and very low standard deviation values, reassuring that the split strategies did not bias our results.

### Using tree-based classifier

For our thorough analysis, we used XGboost [7], which is one of the leading gradient boosting tree-based tools for classification [46]. Differently from deep-learning, XGboost is less computationally expensive and usually does not require a GPU for training, and it can work both with small and large datasets. Additionally, XGboost provides the ability to evaluate and explain the classification rules and rank the features by their importance. We showed that XGboost achieved the best performance over the statistical machine learning algorithms (e.g., SVM and LR) for all datasets. Furthermore, it achieved comparable results to deep learning algorithms that were previously applied on the human dataset *h1*[31, 65].

### Cross-dataset analysis

Most of the previous works trained and tested their predictive models based on a single chimeric miRNA-target dataset (usually *h1*), sometimes complemented by additional experimental data from databases (e.g., [11,66]) or AGO-CLIP data [13,38,49,52,65]. Then these models were evaluated on portions taken out from the training set and in some cases on a few independent datasets from the same or other organisms (Additional file 1, Table S1). The contribution of our work is in providing for the first-time a thorough analysis of all available miRNA-target chimeric datasets, outlining their similarities and dissimilarities. Additionally, we explored the ability to learn classification rules from one dataset and apply them on another dataset, considering all possible combinations of dataset pairs.

The accuracy results of cross-dataset classification ranged between 0.56 to 0.94. To be able to explain these results we examined several factors:

1. Evolutionary distance - we estimated the distance for every pair of organisms (i.e., the time since the organisms diverged from their common ancestor (Table 7)). Among the organisms, the mouse and the human are the closest, with cattle equally and relatively close to them, while *C. elegans* is the most distant from all. Indeed, we got the highest accuracy results when we trained-tested datasets of the same organism and the lowest accuracy results when we trained-tested combinations of *C.elegans* datasets with those from other organisms.
2. Kullback–Leibler (KL) divergence scores - we measured the divergence for every pair of datasets based on their miRNA seed family distribution. Previous analysis of chimeric datasets showed that individual miRNAs exhibit enrichment for specific classes of base-pairing patterns [6,21], suggesting that they may follow different targeting rules. Thus, differences in the distributions of miRNA sequences in the training sets may lead to biases in the rules learned by a machine learning model, which could partially explain the high correlation that we observe between KL-divergence and the classification performance. Interestingly, and maybe not surprisingly, the KL divergence results coincide with the evolutionary distance of the organisms, where *C.elegans* datasets exhibit the highest distance from the datasets of other organisms. The divergence within the same organism is, on average, lower than the divergence between different organisms. This divergence is probably associates with the differences in miRNA distributions among different cell types or developmental stages from which the datasets were generated (Table 1).
3. Area covered in 2D feature space - We visualized the datasets by their features in 2 dimensions using PCA. The visualization highlighted datasets with a lower spread. In particular, *C. elegans* datasets are exceptional relative to the rest of the organisms, concentrated in a narrower area. In addition, the datasets *m1* and *h2*, which represent endogenously ligated chimeras from a mixture of AGO-CLIP experiments, have relatively smaller sizes compared to other datasets from the same organism and have a lower spread. The exception of these two datasets may explain the lower accuracy results obtained in cross-datasets experiments when we used them as training sets.

**Table 7.**
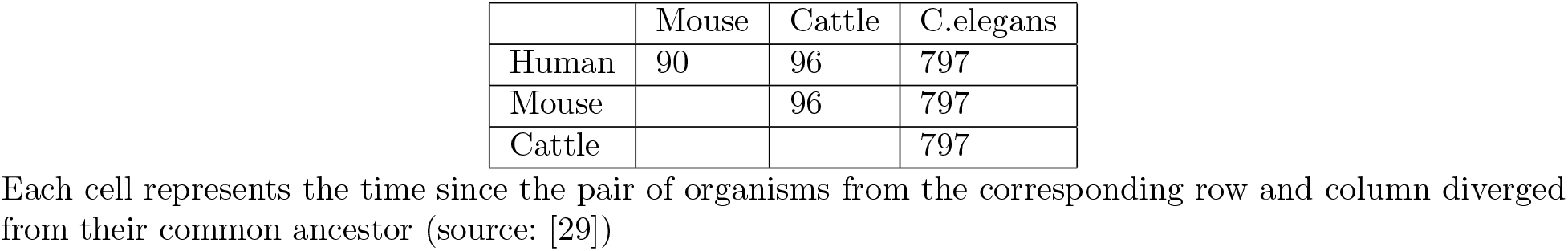
Estimated divergence time [MYA] between organisms in our study

|  | Mouse | Cattle | C.elegans |
| --- | --- | --- | --- |
| Human | 90 | 96 | 797 |
| Mouse |  | 96 | 797 |
| Cattle |  |  | 797 |

Each cell represents the time since the pair of organisms from the corresponding row and column diverged from their common ancestor (source: [29])

## Conclusions

The accuracy results obtained in our cross-datasets experiments are pretty high, as long as the organisms are within a certain evolutionary distance, reflecting the ability of the machine learning model to generalize interaction rules learned from a specific datastet, into more universal interaction rules. Altogether our results suggest that target prediction models could be applied also to organisms where experimental training data is limited or unavailable, as long as they are close enough to the organism that is used for training. As more miRNA-mRNA interaction datasets become available, they could be processed with our pipeline and incorporated into the cross-dataset analysis. Expansion of such analysis on more datasets in the future may also provide insights about the evolution of miRNA-targeting, and identification of general, as well as organism-specific features. Another interesting research direction will be to combine the information from several datasets in an iterative manner and examine the prediction accuracy in close and more distant organisms.

## Materials and methods

### Software packages and tools

Code developed under this research was implemented as a Python package running on a Linux platform. It uses bioinformatics, data analysis and machine learning packages. The bioinformatics packages are ViennaRNA (v2.4.13) [37], Biopython (v1.72) [12] and NCBI Blast [2]. The data packages are pandas (v0.23.4) [42] and numpy (v1.15.4) [47]. The machine learning packages are scikit-learn (v0.20.1) [50] and XGBoost (v0.81) [7].

### Data processing

We acquired eight high-throughput chimeric miRNA-target datasets from four different organisms: human, mouse, cattle, and worm (Table 1). The datasets’ files were downloaded from the journals’ websites [6,19,21,45,55]. In addition, we downloaded miRNA sequences from miRBase (releases 17-22) [27] and 3’UTR sequences from Ensembl Biomart database [58]. Genomic sequences for *C.elegans* were downloaded from wormBase [32], and for human and mouse from UCSC Genome Browser [24]. The datasets were provided in different formats, containing different levels of information about the interactions. Therefore, we developed a processing pipeline to transform the datasets into a standard format, and to include the following fields: metadata (interaction ID, interaction source), miRNA name and sequence, target site sequence (the site where the interaction occurred), and for sites located at the 3’UTRs - the corresponding 3’UTR sequence and the coordinates of the site within it.

We started the pipeline by retrieving the missing miRNA sequences by their name from miRBase (for datasets *ca1, ce2, h3, m2*). Then, we extracted the target sequences (for datasets *ce2, h3, m2*) based on the genomic coordinates. The target sequences are located in various mRNA regions such as 5’ untranslated region (UTR), the coding sequence (CDS), or 3’UTR. miRNA target sites located at the 3’UTRs of mRNA sequences are considered to be most functional [3, 43]. Therefore, in our analyses, we discarded sites that fall outside the 3’UTRs. Since most datasets do not provide the regions containing the interactions, our next step was to obtain that information. We used Blast [2] to match the target mRNA sequences against the 3’UTRs downloaded from the Ensembl Biomart database. We considered only full match results. In cases where multiple UTRs exist per a gene, we considered the longest UTR. The full 3’UTR sequences were kept for the extraction of flanking site features, as later described. Finally, we took the list of miRNA-target pairs and examined the interaction structure. We applied the ViennaRNA suite (RNAduplex) [37] to calculate the interaction duplex, using the miRNA and the target site sequences. We then classified the duplexes based on their seed type: canonical seed, non-canonical seed, and other. Canonical seed interactions have exact Watson-Crick pairing in positions 2–7 or 3–8 of the miRNA, while non-canonical seed interactions may contain *GU* base-pairs and up to one bulged or mismatched nucleotide at these positions [21]. We kept canonical and non-canonical seed interactions only, discarding other interactions from the analysis. Interactions that passed all pipeline stages were designated as positive interactions and were considered for further analysis (Table 8).

**Table 8.** Data processing pipeline

| Paper | [21] | [19] | [55] | [6] | [45] |
| --- | --- | --- | --- | --- | --- |
| Datasets | h1 | ce1, h2, m1 | ca1 | ce2 | h3, m2 |
| miRNA sequence | ✓ | ✓ | mirbase | mirbase | mirbase |
| Target sequence | ✓ | ✓ | ✓ | wormbase | UCSC genome browser |
| Site region | Ensembl Biomart + Blast |  |  |  |  |
| Duplex structure | Vienna RNAduplex |  |  |  |  |
| Seed Filter | Canonical and non-canonical seeds only |  |  |  |  |

The table describes the set of actions required to transform the datasets into a uniform format to serve as input for further data analysis and machine learning experiments. The check-mark sign (✓) represents a piece of information taken directly from the paper without additional calculations.

### Generation of negative interactions

To generate the negative interactions, we used a synthetic method, similarly to the method described in [23, 40, 43]. For each positive interaction appearing in the dataset, we generated a negative interaction as follows. First, we generated a mock miRNA sequence by randomly shuffling the original sequence until there is at most one match in the regions 2-7 and 3-7 between the mock miRNA and any real miRNA of the examined organism (according to miRBase). Next, we provided the mock miRNA and the full 3’UTR sequence as inputs to RNAduplex, which is optimized for computing the hybrid structure between a small probe sequence and a long target sequence. We repeated these two steps until the output duplex had a canonical seed or non-canonical seed. We managed to generate a negative interaction for each positive interaction, thus, at the end of this process, we had balanced datasets.

### Calculation of miRNA distribution

We counted the appearance of each miRNA sequence within a dataset and used that information to generate the cumulative distribution function (CDF) showed in Figure 1. We used the *argmax* function to find the 90% value, which returns the first point in the CDF which is greater than 90%. The seed distribution calculation was done by first clustering miRNA sequences based on their seed sequence (position 2-7) and then following the same steps described above.

### Features

To represent miRNA-target interactions, we used 490 expert-designed features, that are classified into two categories (high-level and low-level) and five subcategories (Table 9). Four of the categories (free energy, mRNA composition, miRNA pairing, and site accessibility) were adopted from [65] while the seed features group was designed in this work. For a full description of the features, see Additional file 4, Table S9.

**Table 9.** Feature categories that are used to represent miRNA-target interactions

| Category | No. of features | Description | Group |
| --- | --- | --- | --- |
| Seed features | 13 | Seed composition and properties | High-level |
| Free Energy | 7 | Free energy of the duplex and the mRNA at different regions | High-level |
| mRNA Composition | 62 | mRNA composition in the site and flanking regions | High-level |
| miRNA Pairing | 38 | Binding information at each miRNA position and across the miRNA-target duplex | Low-level |
| Site accessibility | 370 | Unpaired probabilities of each base | Low-level |
| Total | 490 |  |  |

The *free energy* category includes 7 features representing the minimum free energy of the miRNA-mRNA duplex and the mRNA sequence at different regions including seed, non-seed, site, and flanking regions.

The *mRNA composition* consists of 62 features which supply information about the target mRNA: the distance of the site from the edges of the 3’UTR (2 features), 1- and 2-mer sequence composition within the site region (20 features), and 1- and 2-mer sequence composition of the up and down 70nt flanking region (20 features each).

The *miRNA pairing* category consists of 38 features which describe the duplex itself, including information about base-pairs in each location of the miRNA (20 features), and a total count of base-pairs, mismatches, and gaps in the site region (18 features).

The *site accessibility* features were calculated for each 3’UTR sequence containing the seed site, using RNAplfold in the ViennaRNA package [37] with the following parameters *winsize=80, span=40* and *ulength=10*, as was suggested by previous works [43,65]. The output of RNAplfold provided, for each nucleotide, the mean probability that regions of length 1 to a 10 (ulength), ending at this nucleotide, are unpaired. Out of these calculations we considered only the region that corresponds to the mRNA-target seed region (p2–p8) with 15 flanking bases to either side (37 bases in total), resulting in 37*10=370 features.

In addition to the above features, we designed a new representation for the *seed features*, which describes the base-pairing characteristics of the seed region (nt 1-8 of the miRNA). The new representation includes 13 features: 3 features describe the number of interactions in [nt1-8, nt2-7, nt3-8]; 3 features describe the number of GUs in [nt1-8, nt2-7, nt3-8]; 3 features give information about the number of mismatches (before the first match, inside the seed and after the last match in the seed region); 2 features describe the number of bulges [miRNA side, target side]; and the last two features give additional properties (start with A, index of the first base-pair).

### Splitting the data into training and testing sets

Correct determination of the training and testing sets is crucial for getting reliable results. Specifically, the testing set has to be large enough, it must not contain any sample from the training set, and it has to represent the dataset as a whole. To address these rules, we implemented a stratified random split algorithm. The algorithm ensures that each miRNA appears in the training and the testing sets at the same proportion as in the original dataset. For example, if a specific miRNA constitutes 10% of the interactions in the original dataset, the algorithm ensures that its proportion in both the training and the testing sets is 10%. Within the stratified split, the assignment of the interactions to training (80%) and testing (20%) sets was done randomly according to a random state. For miRNAs appearing once in the dataset, we assigned its interaction to the testing set. We repeated this process 20 times with different random states, yielding 20 training sets and their corresponding 20 testing sets for each dataset. In addition, for each dataset, we generated five control sets by a fully random algorithm, which does not take into account miRNA distributions. We used these sets as a reference baseline, to assess the influence of the stratified split algorithm on the results.

### Evaluation of different machine learning methods

We have chosen six machine learning methods, which are widely used in the field of computational biology, for the classification of miRNA-target interactions: XGBoost [7], Random Forest (RF), K-nearest neighbors vote (KNN), regularized linear models with Stochastic Gradient Descent (SGD), Support Vector Machine (SVM) and Logistic Regression (LR). We performed the following optimization and learning steps for every combination of (dataset, classifier, data split), all together 1200 computationally intensive tasks (Equation 1):

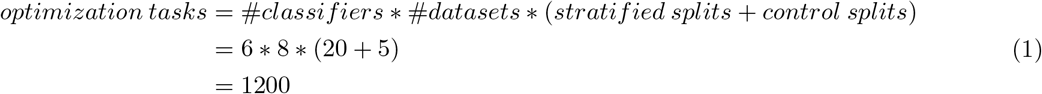

First, we searched for the classifiers’ optimal hyper parameters. We performed an exhaustive search using sklearn GridSearchCV, using 4-fold cross validation, optimized for accuracy performance. Second, we explored the exhaustive search results and identified the set of parameters that achieved the best accuracy results. The classifier corresponding to this set of parameters was saved and used for evaluating the accuracy of classification on the testing set. We provide the values of the parameters for the hyper-parameter optimization in Additional file 3, and accuracy mean and standard deviation for the 20 stratified splits and the 5 control splits in the results section.

We continued with the XGBoost classifier for calculating detailed performance measurements and for the analysis of feature importance. We calculated 6 widely used performance metrics including accuracy (ACC), sensitivity (TPR: true positive rate), and specificity (TNR: true negative rate). In addition, we calculated metrics that are widely used for model comparisons such as the Area Under the Receiver Operating Characteristic Curve (ROC AUC), Matthews Correlation Coefficient (MCC), and F1 score (also known as balanced F-score or F-measure). The equations for the calculation of these metrics are provided in Additional file 1, Equation S1-S5. As before, the mean and the standard deviation for each measure were calculated on all the training-testing splits (Table 5).

### Identification of the top important features

First, we extracted the top features for every dataset as follows. We used the gain metric provided by XGBoost and calculated the mean gain of each feature across the 20 different stratified splits. We then sorted the list of features according to the mean gain. We observed that the top 6 features are the most dominant (for all datasets), and the gain score of the rest is lower in an order of magnitude. Therefore, we kept only the top 6 features of each dataset. Second, to be able to compare between datasets, we scaled the mean gain scores of each dataset to a range of 0-100 (by dividing it by the maximum value and multiplying by 100). Third, we composed a unified list of the top features from all the datasets and generated a table that includes the scaled mean gain values for every feature (row) in every dataset (column). Finally, we calculated the mean score for each feature across all datasets (last column in the table) and sorted the table in descending order to it (see results table 6).

### Calculating Kullback-Leibler divergence

The Kullback-Leibler divergence is calculated on two probability distribution functions and measures the difference and the distance between them according to equation 2.

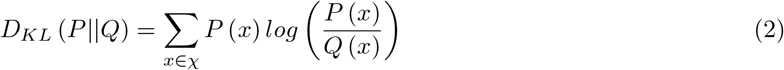

We used the KL divergence to measure the pairwise information loss between every two datasets, which can be interpreted as the amount of information lost when the training set represents the testing set. *P*(*x*) and *Q*(*x*) are the miRNA seed distribution functions as explained in Calculation of miRNA distribution. *Q*(*x*) is the approximation distribution (calculated from the training set) and *P*(*x*) is the true distribution (calculated from the testing set). *χ* is the union of all the miRNA seeds that appear in both datasets.

### Dimensionality reduction using PCA

The dimensionality reduction algorithm enables the representation of the data in a 2-dimensional scatter plot and facilitates the inspection of the data visually. We performed a dimensional reduction using the PCA algorithm to transform the datasets into 2-dimensional representation. We used the same transformer for all datasets to enable their comparison. We extracted the columns corresponding to the top 16 features that were found in section Identification of the top important features (referred to as selected features). Since the datasets are of different sizes, we first oversampled the datasets by random sampler to bring them to the size of the largest dataset. Second, we concatenated the oversampled datasets together. Third, we standardized the selected features by subtracting the mean and scaling to the unit variance for each feature independently. Finally, we fitted a PCA transformer and applied it to the original datasets (without oversampling), yielding the 2D representation of the datasets on the same vector space. The dimensionality reduction was done on the positive experimental interactions only.

### Evaluation of the classification performance between datasets

We next evaluated the performance of XGBoost in the classification of interactions derived from a dataset that is different from the dataset it was trained on. We enumerated over all the 56 possible pairs of training and testing datasets: (*train_i_, test_j_*). For each pair, we loaded 20 XGBoost classifiers (corresponding to 20 splits) generated for dataset *i* in section Evaluation of different machine learning methods and evaluated their performance on the entire dataset *j* (without splitting it). We then computed the mean and the standard deviation of the accuracy results of the 20 tests.

## Declarations

### Ethics approval and consent to participate

Not applicable.

### Consent for publication

Not applicable.

## Author’s contributions

IVL envisioned the project and supervised the work. GBO designed and implemented the processing pipeline and the machine learning system. GBO and IVL planned the evaluation tasks and performed the analysis. GBO and IVL wrote the manuscript. All authors read and approved the final manuscript.

## Acknowledgements

The authors would like to thank DeepMirTar team for providing us the code of their pipeline, which we partially adapted for use in this project.

## Availability of data and materials

The datasets used and/or analysed during the current study are available from the corresponding author on reasonable request.

## Additional Files

### Additional file 1

- Review of Machine-Learning (ML) based methods,
- Supplemental Figure S1 to S2,
- Supplemental Tables S1 to S7,
- Equations S1 to S5.

### Additional file 2

- Table S8. feature importance (XSLX 261KB)

### Additional file 3

- grid_search_params.yaml (YAML 2KB)

### Additional file 4

- Table S9. Features and their definition (XSLX 15KB)

