## Supplementary File S1 for "Comprehensive machine-learning-based analysis of microRNA-target interactions reveals variable transferability of interaction rules across species"

### Supplementary Material

*Gilad Ben Or, Isana Veksler-Lublinsky*

#### 1 REVIEW OF MACHINE-LEARNING (ML) BASED METHODS

Several ML-based methods utilized chimeric miRNA-target datasets to build and evaluate their models. These methods differ in several aspects, including the ML approach and the features used, the choice of datasets for training and testing, inclusion or exclusion of non-canonical interactions from the training/testing sets, and the generation of negative data. Below we provide a summary of some of these methods, focusing on these aspects.

##### 1.1 *Machine learning method and features*

The method chimiRic[8], combines two support vector machine (SVM) classifiers, one to predict miRNA-mRNA duplexes and a second to learn AGO binding preferences. Features used by this method are designed to describe the duplex structure (e.g., binding information at each miRNA position[6], positions of loops' opening and extension), UTR local sequence context, and global positional context.

TarPmiR[3] is a random-forest-based approach that provides a probability that a candidate target site is a true target site. It integrates six conventional features (e.g., folding energy and seed-matching) and seven new features (e.g., largest consecutive pairings).

DeepMirTar[14] is based on a stacked de-noising auto-encoder deep learning method (SdA) and uses 750 different features to describe the interactions (including raw-data-level and expert-designed high-level and low-level features). These features capture the information about the seed match, sequence composition, free energy, site accessibility, conservation, and hot-encoding of miRNAs and their target sites.

miRAW[11] relies on deep artificial neural networks (ANN) to automatically learn the relevant features describing miRNA-gene interactions for predicting miRNA targets. The method works with raw input data and makes no assumptions about suitable input descriptors.

mirLSTM[10] is a deep learning approach that is based on Long Short Term Memory (LSTM). The method captures only the information encoded in the duplex formed between the miRNA and its relevant binding site, which is reduced into a vector over 5 letter alphabet to express four possible base-pairs (*AU,UA,GC,CG*) and one letter for all remaining combinations (including *GU*, bulge, mismatch). These vectors are converted into embedded words to find the relationship between similar sequences using Euclidean distance. Then, the embedded words are fed to the first layer of their LSTM architecture.

MirTarget[13] is based on SVM and uses 50 features that include nucleotide composition, accessibility of the target site, seed conservation, seed base-pairing stability, and location of the target site. In a subsequent study[7], the model was refined with 96 features.

##### 1.2 *Training and testing datasets*

All the above studies trained and tested their models on a dataset of chimeric interactions from human cells generated with the CLASH method[5], referred to as *h1* in the manuscript. In some of the studies, the dataset was filtered based on the location of the sites, seed pairing pattern, or functional evidence. In other studies, it was complemented with additional interactions from other experiments.

DeepMirTar[14] and mirLSTM[10] filtered this dataset to include only canonical and non-canonical sites that are located at the 3'UTRs. They complemented this dataset with an additional small number of interactions retrieved from miRecords[15]). TarPmir[3] used all the interactions available from human CLASH dataset *h1*. chimiRic[8] was trained on a combination of human CLASH dataset and seed-containing sites from AGO-CLIP data. The model was tested on specific miRNA families, by holding out their interactions from the training set. For miRAW[11] the training set was built by intersecting human CLASH, AGO-CLIP and TargetScanHuman[1] datasets with mirTarBase[2] and Diana TarBase[12] to include only validated functional interactions. The initial evaluation was performed on experimentally verified miRNA:gene-targets that were excluded from the training set. miRTarget[13] was trained and tested on human CLASH data combined with chimeras generated by endogenous ligation in human AGO-CLIP experiments[4]. miRTarget v4[7] combined human chimeras with miRNA overexpression data to identify common features that are characteristic of both miRNA binding and target downregulation.

##### 1.3 *Independent testing datasets*

For additional independent testing, the above methods used various datasets, not necessarily derived from ligation-based experiments. DeepMirTar[14] was tested on human PAR-CLIP dataset. mirLSTM[10] was tested on a small experimental dataset. TarPmir[3] was tested on two PAR-CLIP human datasets and a HITS-CLIP dataset from mouse. chimiRic's[8] model was tested on chimeric interactions from *C. elegans*[4] and mouse[9]. miRAW[11] performed additional evaluations on microarray datasets reporting mRNA changes after transfecting a miRNA into HeLa cells. miRTarget[13] and miRTarget v4[7] tested their model on mRNA microarray datasets from miRNA knockdown experiments. miRTarget v4 performed additional testing on HITS-CLIP data from the mouse brain.

##### 1.4 *Negative datasets*

For negative datasets, DeepMirTar[14] and mirLSTM[10] used mock miRNAs to generate a negative interaction for each positive one. TarPmir[3] generated negative target sites on positive mRNAs, such that they do not overlap with positive sites and have similar nucleotide composition compared to the positive sites. miRAW built a negative dataset based upon experimentally verified data. chimiRic[8] generated negative set from canonical miRNA seed matches that are not AGO bound based on CLIP data, together with (miRNA, site) pairs where an AGO-bound site is paired with an incorrect miRNA based on CLASH chimeras. miRTarget[13] and miRTarget v4[7] generated negative examples based on CLIP data, selecting non-target sites based on a set of criteria, e.g., no overlap with a positive site, detectable expression of the transcript based on microarrays, and the existence of perfect seed match to one of the miRNAs expressed in the cells.

**Figure S1.** Classification of the negative miRNA-target duplexes based on their base-pairing patterns

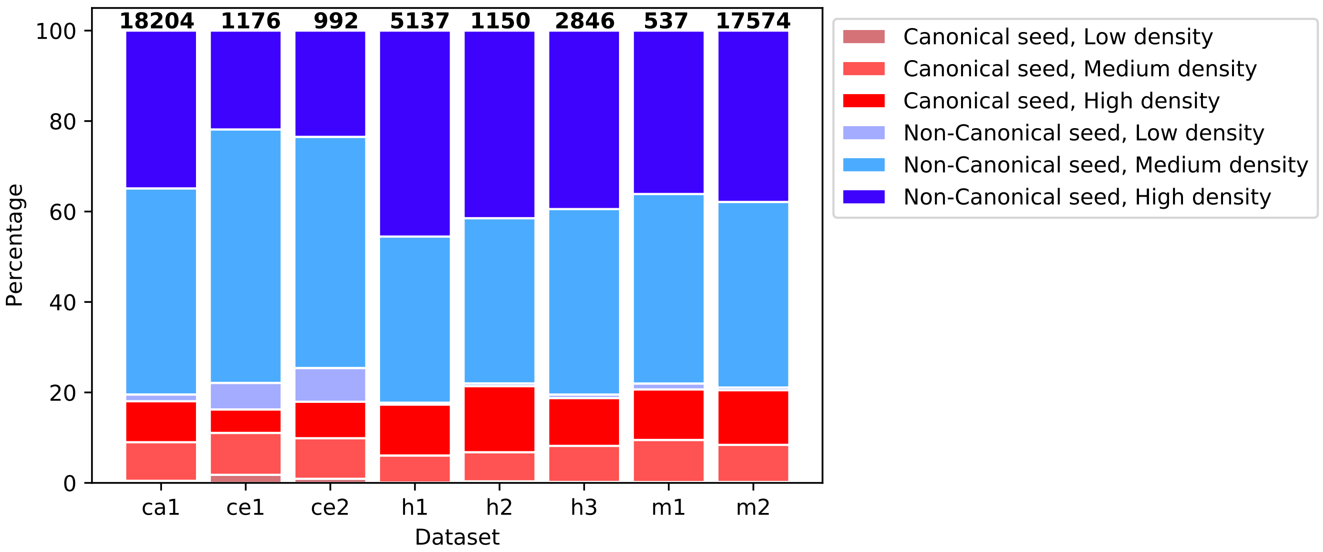


Distribution of miRNA-target duplexes across 6 classes according to the seed type (canonical and non-canonical) and to the base-pairing density (low: less than 11bp, medium: 11-16bp and high: more than 16bp).

**Figure S2.** Results of cross-dataset classification with 16 features

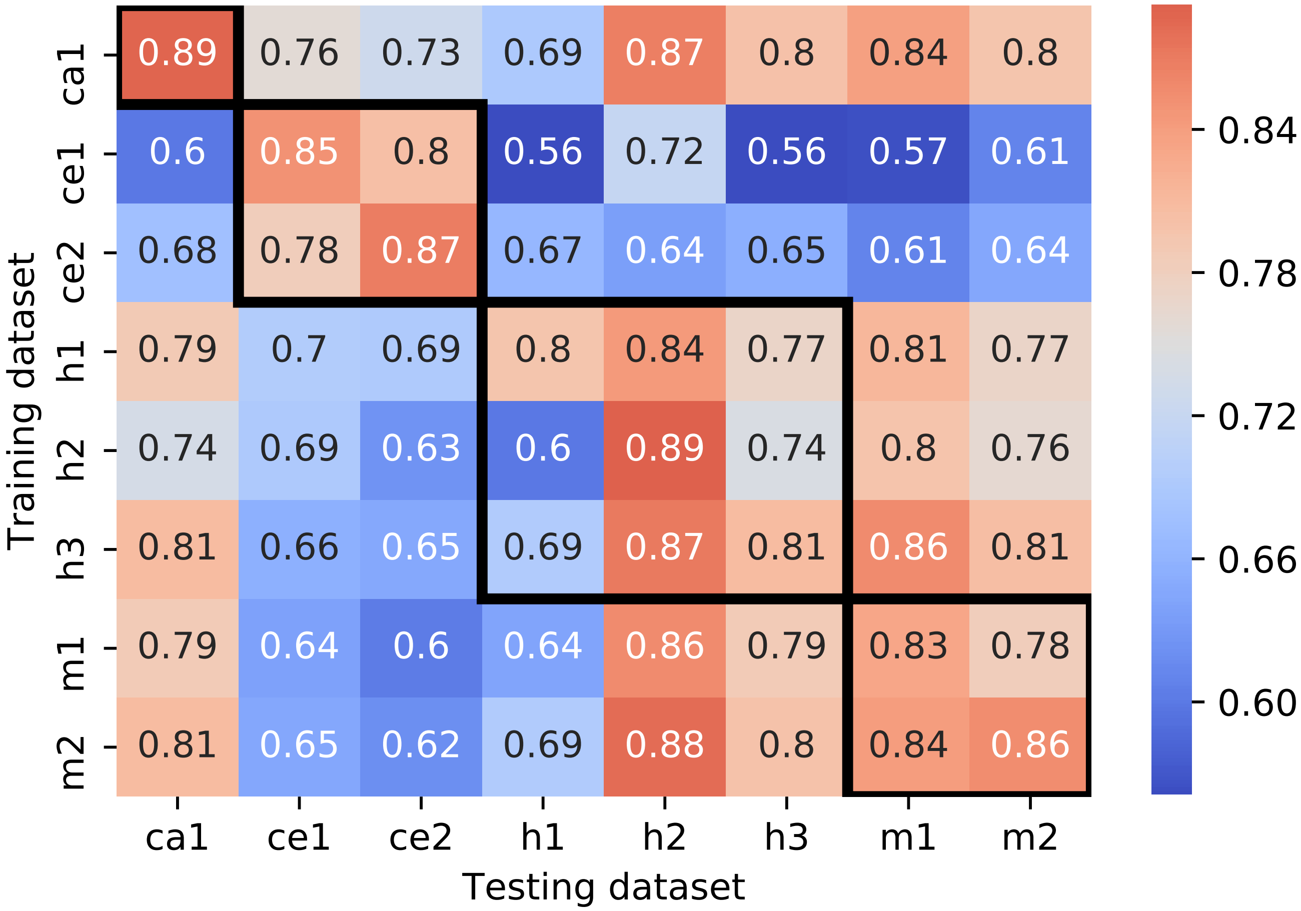


Each cell *(i,j)* represents the mean accuracy of the 20 classifiers that were trained on 20 training-testing dataset splits of dataset *i* and tested on dataset *j* *ACC(i, j)* using 16 features found in section *Top important features of each dataset*. The black frames surround the results of dataset pairs originating from the same organism.

**Table S1.** A summary of machine-learning based methods that utilized chimeric miRNA-target datasets in their models.

| Tool | ML method | Datasets used for training/testing | Independent Dataset | Negative interactions | Features |
|---|---|---|---|---|---|
| chimiRic[8] | SVM | Human (CLASH, AGO-CLIP) | chimeras from Mouse and *C.elegans* | seed matching non-CLIP sites | Small |
| TarPmiR[3] | Random Forest | Human (CLASH) | Human (PAR-CLIP), Mouse (HITS-CLIP) | Negative target sites, on positive mRNAs | 13 |
| DeepMirTar[14] | Deep learning | Human (CLASH) + mirRecords | Human (PAR-CLIP) | Mock miRNAs | 750 |
| miRAW[11] | Deep learning | Human (intersection of: CLASH, CLIP, TargetScan with Diana TarBase, mirTarBase) | Human (microarray) | Experimental data | Raw sequence |
| mirLSTM[10] | Deep learning | Human (CLASH) + mirRecords | Experimental | Mock miRNAs | Raw sequence |
| mirTarget[13] | SVM | Human (CLASH, AGO-CLIP) | Human(microarrays) | seed matching non-CLIP sites on expressed mRNAs | 50 |
| mirTarget v4[7] | SVM | Human (Intersection of CLASH and microarrays) | Mouse (HITS-CLIP), Human (microarrays) | seed matching non-CLIP sites on expressed mRNAs | 96 |

**Table S2.** The accuracy performance of different tools (left) and machine learning methods (right), on human dataset referred to as *h1* in the manuscript, taken from deepMirTar [14].

| Methods | ACC |
|---|---|
| Miranda | 0.6592 |
| RNAhybrid | 0.6988 |
| PITA | 0.4981 |
| TargetScan v7.0a | 0.5801 |
| TarPmiR | 0.7446 |
| DeepMirTar (SdA) | 0.9348 |

| Machine learning alg. | ACC |
|---|---|
| DT (Decision Tree) | 0.8139 (0.0137) |
| BNB (Bernoulli Naïve Bayes) | 0.7570 (0.0098) |
| LR (Logistic Regression) | 0.8491 (0.0117) |
| RF (Random Forest) | 0.8811 (0.0090) |
| MLP (Multi-Layer Perceptron) | 0.8990 (0.0099) |
| CNN-1D (Convolutional Neural Network) | 0.8886 (0.0145) |
| CNN-2D (Convolutional Neural Network) | 0.8765 (0.0169) |

The right table contains the mean and the standard deviation values (in brackets) acquired from 20 models that were built based on different random splits of the dataset to training/validation/testing.

**Table S3.** Classification accuracy of xGBoost when applied to random (control) training-testing dataset splits

| Dataset | Accuracy |
|---|---|
| ca1 | 0.938 (0.004) |
| ce1 | 0.891 (0.013) |
| ce2 | 0.89 0(0.012) |
| h1 | 0.832 (0.006) |
| h2 | 0.920 (0.013) |
| h3 | 0.849 (0.008) |
| m1 | 0.848 (0.011) |
| m2 | 0.897 (0.006) |

The cells contain the mean and the standard deviation values (in brackets) acquired from 5 models that were trained and evaluated on different random dataset splits.

**Table S4.** Intra-dataset classification accuracy of xGBoost classifier trained on an extended 580 features set (including one-hot encoding features)

| Dataset | Accuracy |
|---|---|
| ca1 | 0.985 (0.001) |
| ce1 | 0.951 (0.006) |
| ce2 | 0.959 (0.008) |
| h1 | 0.93 (0.006) |
| h2 | 0.922 (0.008) |
| h2 | 0.908 (0.007) |
| m1 | 0.877 (0.014) |
| m2 | 0.956 (0.002) |

The cells contain the mean and the standard deviation (in brackets) values acquired from 20 models that were trained and evaluated on different training-testing stratified dataset splits.

**Table S5.** Feature importance

| Feature/Dataset | ca1 | ce1 | ce2 | h1 | h2 | h3 | m1 | m2 | mean |
|---|---|---|---|---|---|---|---|---|---|
| **Number of GU bp within the seed** | 225 | 25 | 51 | 25 | 23 | 77 | 12 | 172 | 76 |
| **bp in the 1st nt of the seed** | 142 | 23 | 18 | 63 | 14 | 23 | 11 | 147 | 55 |
| Number of GU bp within the site | 94 | 20 | 17 | 89 | 11 | 41 | 15 | 47 | 42 |
| **Number of bp at location 2-7** | 95 | 9 | 53 | 11 | 10 | 28 | 6 | 31 | 30 |
| Duplex minimum free energy | 29 | 13 | 6 | 9 | 58 | 14 | 15 | 90 | 29 |
| Proportion of G in mRNA at the site region | 26 | 21 | 6 | 10 | 21 | 25 | 42 | 64 | 27 |
| Proportion of GG in mRNA at the site region | 67 | 6 | 5 | 11 | 4 | 23 | 33 | 45 | 24 |
| **bp in the 4th nt of the seed** | 19 | 29 | 11 | 9 | 6 | 12 | 1 | 21 | 13 |
| **bp in the 5th nt of the seed** | 27 | 8 | 8 | 13 | 3 | 12 | 12 | 20 | 13 |
| Number of bulges outside the seed | 7 | 17 | 3 | 22 | 18 | 7 | 4 | 14 | 12 |
| **Number of GC bp within the seed** | 15 | 6 | 13 | 16 | 7 | 10 | 4 | 20 | 11 |
| **bp in the 2nd nt of the seed** | 18 | 12 | 20 | 6 | 6 | 10 | 6 | 10 | 11 |
| Accessibility (nt=21, len=10) | 19 | 5 | 4 | 6 | 14 | 6 | 5 | 11 | 9 |
| minimum free energy of the target site + 50nt flanking regions | 18 | 3 | 3 | 6 | 5 | 8 | 15 | 10 | 8 |
| Number of GC bp outside the seed | 8 | 8 | 6 | 9 | 16 | 6 | 3 | 8 | 8 |
| **Number of mismatches inside the seed** | 10 | 1 | 8 | 17 | 0 | 10 | 1 | 16 | 8 |

The table shows 16 features representing the union of the top 6 features of each dataset, along with their gain values which were computed by XGBoost. The features are ordered by their mean gain across all datasets. This is an unscaled version of Table 6.

**Table S6.** Cross-dataset classification results (complementary information to Figure 6)

| | ca1 | ce1 | ce2 | h1 | h2 | h3 | m1 | m2 |
|---|---|---|---|---|---|---|---|---|
| ca1 | 0.937 (0.002) | 0.779 (0.005) | 0.733 (0.01) | 0.699 (0.003) | 0.902 (0.002) | 0.839 (0.002) | 0.885 (0.006) | 0.823 (0.002) |
| ce1 | 0.61 (0.005) | 0.889 (0.014) | 0.809 (0.011) | 0.561 (0.006) | 0.732 (0.009) | 0.575 (0.005) | 0.598 (0.008) | 0.626 (0.006) |
| ce2 | 0.688 (0.008) | 0.806 (0.007) | 0.891 (0.016) | 0.654 (0.008) | 0.647 (0.008) | 0.667 (0.008) | 0.643 (0.01) | 0.657 (0.006) |
| h1 | 0.794 (0.005) | 0.714 (0.007) | 0.686 (0.011) | 0.824 (0.007) | 0.864 (0.006) | 0.792 (0.004) | 0.815 (0.008) | 0.786 (0.004) |
| h2 | 0.725 (0.004) | 0.712 (0.005) | 0.628 (0.007) | 0.582 (0.004) | 0.904 (0.007) | 0.727 (0.004) | 0.78 (0.008) | 0.753 (0.005) |
| h3 | 0.832 (0.004) | 0.691 (0.005) | 0.654 (0.009) | 0.693 (0.006) | 0.898 (0.003) | 0.835 (0.007) | 0.872 (0.007) | 0.828 (0.003) |
| m1 | 0.782 (0.007) | 0.638 (0.01) | 0.591 (0.011) | 0.623 (0.006) | 0.853 (0.006) | 0.771 (0.005) | 0.847 (0.015) | 0.778 (0.005) |
| m2 | 0.856 (0.002) | 0.705 (0.004) | 0.663 (0.005) | 0.702 (0.002) | 0.908 (0.003) | 0.83 (0.003) | 0.891 (0.004) | 0.9 (0.004) |

Each cell *(i,j)* represents the mean accuracy and standard deviation of the 20 classifiers that were trained on dataset *i* and tested on dataset *j* *ACC(i, j)*.

**Table S7.** Intra-dataset classification accuracy with 16 features only

| Dataset | XGBoost | RF | KNN | SGD | SVM | LR |
|---|---|---|---|---|---|---|
| ca1 | 0.889 (0.004) | 0.889 (0.003) | 0.835 (0.003) | 0.759 (0.013) | 0.842 (0.003) | 0.771 (0.005) |
| ce1 | 0.851 (0.015) | 0.85 (0.014) | 0.782 (0.015) | 0.778 (0.019) | 0.829 (0.015) | 0.794 (0.017) |
| ce2 | 0.87 (0.017) | 0.868 (0.017) | 0.809 (0.014) | 0.817 (0.032) | 0.859 (0.016) | 0.837 (0.015) |
| h1 | 0.796 (0.007) | 0.792 (0.006) | 0.748 (0.008) | 0.733 (0.011) | 0.784 (0.008) | 0.743 (0.008) |
| h2 | 0.892 (0.012) | 0.885 (0.008) | 0.852 (0.009) | 0.861 (0.021) | 0.884 (0.009) | 0.881 (0.008) |
| h3 | 0.809 (0.009) | 0.805 (0.007) | 0.761 (0.011) | 0.711 (0.047) | 0.804 (0.004) | 0.754 (0.007) |
| m1 | 0.832 (0.013) | 0.833 (0.015) | 0.746 (0.018) | 0.71 (0.071) | 0.808 (0.014) | 0.762 (0.016) |
| m2 | 0.855 (0.004) | 0.848 (0.004) | 0.808 (0.003) | 0.785 (0.015) | 0.845 (0.004) | 0.794 (0.004) |

Intra-dataset classification accuracy performed with different machine learning methods that incorporate only 16 features identified in section *Top important features of each dataset*. The cells contain the mean and the standard deviation (in brackets) values of the accuracy results acquired from 20 models that were trained and evaluated on different training-testing dataset splits.

$$ACC = \frac{TP + TN}{TP + FP + FN + TN}$$ **Equation S1.**

$$TPR = \frac{TP}{TP + FN}$$ **Equation S2.**

$$TNR = \frac{TN}{TN + FP}$$ **Equation S3.**

$$MCC = \frac{TP * TN - FP * FN}{\sqrt{(TP + FP)(TP + FN)(TN + FP)(TN + FN)}}$$ **Equation S4.**

$$F1score = \frac{TP}{TP + \frac{1}{2}(FP + FN)}$$ **Equation S5.**
